# Nuclear force transmission drives cancer-associated fibroblast activation under BRAF inhibition

**DOI:** 10.1101/2025.07.19.665630

**Authors:** Jie Wang, Bruna da Silva Soley, Yao Xiao, Lindsey G. Siegfried, Linli Zhou, Mingang Xu, Sarah E. Millar, Thomas Andl, Yuhang Zhang

## Abstract

Cancer-associated fibroblasts (CAFs) exhibit striking plasticity, enabling them to adapt to external cues such as therapeutic and mechanical stress in the tumor microenvironment (TME). Here, we uncover a shared mechanotransduction pathway through which both BRAF inhibition and matrix stiffness converge in nuclear remodeling to drive CAF activation. Mechanistically, BRAF inhibitors (BRAFis) accelerate RAS-dependent RAF homo and heterodimerization and ERK signaling, leading to GSK-3β inactivation and Rho kinase (ROCK) activation. In parallel, stiff substrates engage integrin receptors to directly activate ROCK signaling. Activation of ROCK induces actin stress fiber assembly, generating mechanical forces that deform the nucleus. In both contexts, nuclear reshaping promotes β-catenin translocation and actin polymerization through a feedback loop that continuously enforces CAF activation and promotes melanoma progression. Notably, pharmacological inhibition of ROCK activity blocks both BRAFi- and stiffness-induced nuclear remodeling and β-catenin accumulation, identifying the ROCK–cytoskeleton–nucleus axis as a critical regulator of CAF adaptation and functionality. Collectively, these findings reveal a mechanically tuned nuclear signaling hub that integrates chemical and physical cues to promote stromal adaptation and suggest ROCK inhibition as a strategy to counteract therapy-induced fibroblast reprogramming and improve therapy response.

## INTRODUCTION

Cancer-associated fibroblasts (CAFs) are a dominant and functionally versatile stromal population in the tumor microenvironment (TME), orchestrating critical processes that underpin tumor progression, metastatic dissemination, and therapy resistance^1^. Arising from diverse cellular origins, including resident fibroblasts, mesenchymal stem cells, or other progenitor cells, CAFs exhibit a reprogrammed phenotype marked by the expression of specific biomarkers, such as α-smooth muscle actin (α-SMA), fibroblast activation protein (FAP), and platelet-derived growth factor receptor β (PDGFR-β)^2–4^. Functionally, CAFs contribute to tumor progression by remodeling the extracellular matrix (ECM), secreting pro-tumorigenic cytokines and growth factors, and engaging in dynamic crosstalk with malignant tumor and stromal cells, thereby shaping the biochemical and mechanical landscape of the TME^4^.

Within the TME, CAFs encounter and integrate a wide spectrum of biophysical and biochemical signals, including therapeutic agents, ECM stiffness, and metabolic byproducts, which, in turn, modulate their activation state, phenotype, and function. Notably, mechanical stress resulting from matrix stiffening and ECM remodeling activates mechanotransduction pathways in CAFs, including integrin signaling and YAP/TAZ nuclear translocation, which increase cytoskeletal tension and further reinforce CAF activation^5,6^. In this activated state, CAFs secrete pro-survival and immunosuppressive factors and contribute to a fibrotic ECM that impedes drug delivery and fosters therapeutic resistance^4^. Equally important, therapeutic agents, ranging from cytotoxic to targeted therapies, represent dynamic components of the TME and are emerging as active participants in remodeling the TME and exerting selective pressure on both tumor and stromal cells^7^. However, the convergence of chemical and physical cues in regulating CAF behavior remains poorly understood.

One of the most transformative advances in the treatment of metastatic melanoma has been the development of BRAF inhibitors (BRAFis) targeting the V600E mutation in the BRAF kinase domain^8^. This mutation allows BRAF to bypass autoinhibition and constitutively activate the downstream MAPK/ERK pathway, driving tumor growth^9–12^. BRAFis, such as vemurafenib and dabrafenib, effectively block this aberrant signaling; however, therapeutic resistance frequently emerges, limiting long-term patient benefits^13,14^. While BRAFis were designed to selectively target mutant cancer cells, their off-target effects on stromal cells, including CAFs that carry wild-type BRAF, are increasingly recognized as critical factors influencing therapeutic success and resistance^15,16^. Growing evidence has revealed paradoxical activation of the MAPK pathway in wild-type BRAF or RAS-mutant cells via BRAFi-induced RAF dimerization^17–19^. Despite these findings, the mechanisms by which BRAFis modulate CAFs and their potential interactions with mechanical signaling pathways remain unresolved.

In this study, we sought to delineate the molecular basis by which external cues, both pharmacological and mechanical, drive adaptive reprogramming of CAFs in melanoma. We showed that BRAFis recapitulate mechanical stress responses in CAFs by activating the ROCK– MYPT1 signaling axis, which promotes actin stress fiber assembly and nuclear elongation. This cytoskeletal-to-nuclear mechanical relay facilitates β-catenin nuclear translocation, establishing β-catenin as a common effector of pharmacological and mechanical inputs. Intriguingly, our results highlight a unified actin-β-catenin feedback mechanism that integrates chemical and mechanical cues to regulate CAF adaptation and activation. Furthermore, inhibition of ROCK signaling abrogates BRAFi- and matrix stiffness–induced nuclear deformation and β-catenin activation, underscoring the ROCK–cytoskeleton–nucleus axis as a key vulnerability in the CAF-driven stromal response. Together, these results establish actin-mediated nuclear architecture as a master regulator of CAF function and reveal a therapeutic opportunity to disable stromal adaptability and enhance the efficacy of targeted therapies in melanoma.

## MATERIALS AND METHODS

### Transgenic mouse strains

*Col1α2-CreER* mice were kindly provided by Dr. de Crombrugghe^20^. The *tetO-ΔN-β-catenin* transgenic mouse strain was generated in Dr. Sarah Millar’s laboratory and was reported in our previous studies^21,22^. The *Rosa-rtTA* (Jax 005670) mouse strain was obtained from the Jackson Laboratory (Bar Harbor, ME, USA). Both mouse strains were backcrossed to the C57BL/6 (B6) genetic background. To generate the triple *Col1α2-CreER*; *Rosa-rtTA*; *tetO-ΔN-β-catenin* mouse strain, *Col1α2-CreER*, *Rosa-rtTA*, and *tetO-Δ N-β-catenin* mice were crossed for several generations. Mice were genotyped by polymerase chain reaction (PCR) analysis of genomic DNA extracted from tail biopsies. The presence of the Cre transgene, *Col1α2-CreER*, was identified by PCR using the forward primer, 5′ CGGTCTGGCAGTAAAAACTAT 3′, and the reverse primer, 5′ CAGGGTGTTATAAGCAATCCC 3′. The *Rosa-rtTA* allele was genotyped using the forward primer, 5 ′ AAGACCGCGAAGAGTTTGTC 3 ′ , and the reverse primer, 5 ′ AAAGTCGCTCTGAGTTGTTAT 3 ′ . The *ΔN-β-catenin* transgene was genotyped using the forward primer, 5 ′ CCTTGTATCACCATGGACCCTCAT 3 ′ , and the reverse primer: 5 ′ TAGTGGGATGAGCAGCGTCAAACT 3′. Standard PCR protocols were used. Male and female animals were used in the study, and no differences were observed between sexes. All mice were housed in the Laboratory Animal Services Facility of the University of Cincinnati under an artificial 14 hour/10 hours light-dark cycle and allowed free access to food and water. The Institutional Animal Care and Use Committee of the University of Cincinnati (IACUC) approved all experimental procedures involving mice (22-08-19-01).

### Cell lines

Human melanoma cell lines, A375 (Cat# CRL-1619) and SK-MEL-24 (Cat# HTB-71), were purchased from the American Type Culture Collection (ATCC, Manassas, VA). Two human CAF cell lines, 224350P1 and DT01027P1, isolated from surgically excised human melanoma tissues, were obtained from Asterand Bioscience (Detroit, MI) and designated as M50 and M27, respectively. Human dermal fibroblasts were provided by Dr. Dorothy Supp of the Department of Surgery at the University of Cincinnati. Murine *Braf^V600E^; Pten^lox5/lox5^* D4M melanoma cells were purchased from Kerafast (Boston, MA)^23^. Melanoma cell lines were authenticated by ATCC and routinely validated in the laboratory based on the expression of melanoma proteoglycan antigens and surface molecules, including GP100^24^, MITF, melan-A, and tyrosinase^25^. We authenticated CAFs and normal human dermal fibroblasts by evaluating their unique morphological characteristics and expression of specific skin fibroblast markers, including TE7^26^, vimentin, and α-SMA^27^. All cell lines were passaged fewer than 20 times and maintained in Dulbecco’s modified Eagle medium (DMEM) supplemented with 10% (v/v) fetal bovine serum (FBS), 1% (v/v) 10,000 units/mL penicillin, 10,000 μg/mL streptomycin, and 25 μg/mL Amphotericin B in a humidified incubator at 37°C with 5% CO2. The cells were routinely monitored for mycoplasma contamination using a Universal Mycoplasma Detection Kit. Cell culture reagents were purchased from Corning Life Sciences (Durham, NC) unless otherwise stated. Experimental procedures involving biosafety hazards were performed under the University of Cincinnati Institutional Biosafety Committee protocol 16-08-17-01.

### Mouse primary fibroblast isolation

The sex of neonatal *Col1α2-CreER*, *Rosa-rtTA*, and *TetO-ΔN-β-catenin* mice was first determined by genotyping the SRY gene using PCR with the forward primer, 5’ TTGTCTAGAGAGCATGGAGGGCCATGTCAA 3’, and the reverse primer, 5’ CCACTCCTCTGTGACACTTTAGCCCTCCGA 3’. To isolate dermal fibroblasts, the dorsal skin of sex-matched 2–3-day-old littermates was collected, and subcutaneous fat and other tissues were removed from the dermal side. The skin was floated with the epidermis facing up in a 0.25% trypsin solution at 37°C for one hour to separate the dermis and epidermis. The dermis was minced into small pieces of approximately 1 mm^2^ using sterile scissors. The dermal pieces were then incubated in 5 mL of 2.5 mg/mL collagenase IV solution for 30 minutes at 37°C with pipetting every 10 minutes. After 30 minutes, 5 mL of DMEM was added to stop the digestion. The dermal cell suspension was filtered through 70 μm and 40 μm strainers to remove cell clumps and undissociated tissue. Fibroblasts were collected by centrifugation at 1500 rpm for five minutes and resuspended in 10 mL of DMEM for culture in a 10 cm dish. Fibroblast genotypes were determined by PCR as described for transgenic mouse strains and validated by immunostaining for the expression of α-SMA, vimentin, S100A4, keratin 14 (negative, keratinocyte marker), and TRP1 (negative, melanocyte marker). Four pairs of ΔN-β-catenin-overexpressing mouse fibroblasts and wild-type fibroblasts were isolated and frozen in liquid nitrogen at passages 3-5 for *in vitro* and *in vivo* studies.

### Induction of ΔN-β-catenin expression in cultured mouse fibroblasts

To activate CreER- and rtTA-dependent ΔN-β-catenin expression in isolated mouse fibroblasts, 3 × 10^5^ cells were seeded in a 10 cm dish and treated with 4-hydroxytamoxifen (4-OHT; Sigma‒ Aldrich, Cat# H6278) at a final concentration of 0.2 μg/mL and doxycycline (Dox; Sigma‒Aldrich, Cat# 33429) at 1.25 μg/mL for 72 hours. Following induction, cells were washed with PBS and harvested for subsequent assays.

### Cell proliferation assay

1 × 10^5^ Dox-treated mouse fibroblasts of the genotypes *Col1α2-CreER; Rosa-rtTA*; *TetO-ΔN-β-catenin* or *Col1α2-CreER; Rosa-rtTA* were seeded and cultured in a 6-cm dish. On days three, five, and seven, the cells were collected, and the number of cells was counted via a hemocytometer. At least three repeats were included for each indicated cell line. Cell number counting was performed at least three times for statistical analysis.

### Collagen gel contraction assay

Dox-treated 5 × 10^4^ mouse fibroblasts of the genotypes *Col1α2-CreER; Rosa-rtTA*; *TetO-ΔN-β-catenin* and *Col1α2-CreER*; *Rosa-rtTA* were suspended in 500 µL of 1 mg/mL rat tail collagen type I in DMEM and seeded in one well of a 24-well tissue culture plate. After a 30-minute incubation in a humidified incubator at 37°C, 1 mL of fresh medium was added to the top of the gel for incubation. After a 72-hour incubation, the gel was detached from the wall of each well and allowed to contract as indicated. Images of the gels were taken using a Nikon digital camera every 24 hours, and ImageJ software (NIH, Bethesda, MD) was used to measure the area of the gel. Gel contraction was calculated as the percentage reduction in the gel area from 0 to 72 hours: gel contraction (%) = [(area at 0 h − area at 72 h)/area at 0 h] × 100.

### Confocal reflection microscopy (CRM)

5 × 10^4^ Dox-treated mouse fibroblasts of the genotypes *Col1α2-CreER; Rosa-rtTA*; *TetO-ΔN-β-catenin* or *Col1α2-CreER*; *Rosa-rtTA* were embedded in collagen gels in the same way as described for the collagen gel contraction assay. After a 72-hour incubation, the gels were collected, and collagen fiber distribution and organization were directly examined using a Zeiss LSM 710 confocal microscope at 40X magnification. Images were acquired at a depth of at least 100 µm below the top surface of the gel to minimize the edge effects. Images of at least ten areas of each gel were randomly captured for 3D reconstruction using the ImageJ software. Fiber connectivity and mean spacing were calculated using ImageJ with the BoneJ plugin (http://bonej.org).

### Mouse melanoma induction

As shown in Fig. 4A, two groups of melanomas were generated by intradermal injection of a mixture of 1 × 10^5^ D4M melanoma cells and 1 × 10^5^ uninduced mouse fibroblasts of the genotype *Col1α2-CreER; Rosa-rtTA*; *TetO-ΔN-β-catenin* or *Col1α2-CreER; Rosa-rtTA* into the flanks of 5-week-old B6 mice. For each group, a minimum of three male and three female mice were used as recipient mice. The mice were monitored daily after cell injection for tumor formation. After the tumors reached a volume of approximately 62.5 mm^3^ (marked as day zero), to induce β-catenin overexpression in *Col1α2-CreER; Rosa-rtTA*; *TetO-ΔN-β-catenin* fibroblasts, all mice were fed a Dox diet (Bio-Serv, F-4096) and underwent intraperitoneal injection of 10 mg/mL tamoxifen (Sigma‒Aldrich, St. Louis, MO) in corn oil at 1 mg/g body weight for seven consecutive days. Afterwards, the tumor size was measured using a caliper every other day for 18 days. Tumor volume was calculated using the following formula: [(width)^2^ × (length)]/2. The mice were euthanized around day 18 when the tumor size exceeded 15% of the body size, and the tumors were harvested for analysis. The number of mice used in each study was calculated using a power analysis. Specific randomization and blinding were not necessary in the reported study. Student’s t-test was used to determine the significance of differences in tumor size and weight between the two groups.

### Histology, immunofluorescence staining, and immunohistochemistry

The collected melanoma tissues were fixed in 10% formalin overnight at 4°C and embedded in paraffin for sectioning. Five μm-thick paraffin-embedded tumor tissue sections were prepared for hematoxylin and eosin (H&E) staining and immunostaining as described previously^7,28^. For histological analysis, paraffin sections were stained using a standard H&E staining protocol, mounted with VectaMount permanent mounting medium (Vector Labs, Burlingame, CA), and analyzed using a Nikon Eclipse 80i fluorescence microscope.

For immunostaining, paraffin sections were deparaffinized, rehydrated, and unmasked in citrate buffer (pH 6.0) using the microwave heating method. After washing with PBS, the sections were blocked with 10% bovine serum albumin (BSA) and subsequently incubated with each primary antibody at 4°C overnight. The following primary antibodies were used: anti-α-SMA (Thermo Fisher, clone 1A4, 1:200), anti-fibronectin (Millipore, clone F3648, 1:200), anti-Ki67 (Thermo Fisher, clone 27309-1-AP, 1:50), and anti-cyclin D1 (Thermo Fisher, clone SP4, 1:50). After incubation with primary antibodies, the slides were washed with PBS three times, incubated with the corresponding biotin-conjugated secondary antibodies at room temperature for one hour, washed again three times with PBS, and then incubated with either fluorochrome-conjugated streptavidin for immunofluorescence or VECTASTAIN Elite ABC Reagents (Vector Laboratories, Burlingame, CA) for immunohistochemistry. Nuclei were counterstained with DAPI (blue) for immunofluorescence and hematoxylin (blue) for immunohistochemistry. Images were captured using a Nikon Eclipse 80i fluorescence microscope.

The numbers of positively stained cells (α-SMA+, Ki67+, Cyclin D1+) or α-SMA-melanoma cells in each high-power field (40X) were counted using the particle analysis function of ImageJ. The number of cells per square millimeter was calculated by multiplying the number of cells counted in each field by 4.5. The fibronectin+ area in each high-power field (40X) was measured using the ImageJ software. The fibronectin+ (%) was calculated by dividing the fibronectin+ area by the area of the entire field. For each melanoma sample. At least 12 randomly selected fields were counted for each sample. A minimum of three independent experiments were performed for each quantification of immunostaining.

### Collagen staining

Picrosirius red staining was performed using the Picro-Sirius Red Stain Kit according to the manufacturer’s instructions (American MasterTech, Lodi, CA). Briefly, paraffin sections of mouse melanomas were dewaxed, hydrated, and stained with picrosirius red for one hour. After staining, the tissue sections were washed with 1% acetic acid for one minute, dehydrated in 100% ethanol, cleared in xylene, and mounted using VectaMount permanent mounting medium (Vector Laboratories) for evaluation.

### Quantification of collagen content

Paraffin sections were deparaffinized and rehydrated for collagen content quantification using the Sirius Red/Fast Green Collagen Staining Kit (Chondrex, Cat# 9046) following the manufacturer’s instructions. Briefly, after the slide was incubated with the dye solution at room temperature for 30 minutes, the dye solution was removed, and the slide was rinsed with distilled water until the water became colorless. One milliliter of dye extraction buffer was added to each slide to elute the dye from the dyed tissues. Optical absorbances at 540 nm and 605 nm was measured using a microplate reader. The collagen content was calculated using the following formula: collagen (μg/section) = [OD540 − (OD605 × 0.291)]/0.0378. Three tissue sections per group were used for quantification and statistical analyses.

### Generation of transgene-expressing human CAF lines

To immortalize M27 and M50 cells, Lenti-SV40 lentivirus (ABM, Cat# G203) was purchased from Applied Biological Materials (Richmond, Canada). The immortalized cell lines were named iM27 and iM50. All BRAF and CRAF constructs, including Flag-tagged BRAF (BRAF_OHu28570C_pGenlenti_N-FLAG), Myc-tagged BRAF (HygR_BRAF_OHu28570C_pGenlenti_N-MYC), BRAF-T529N (BRAF-T529N_OHu28570C_pGenlenti_N-FLAG), Flag-tagged CRAF (HygR_RAF1_OHu28584C_pGenlenti_N-Flag), Myc-tagged CRAF (RAF1_OHu28584C_pGenlenti_N-MYC), and CRAF-T421N (RAF1-T421N_OHu28584C_pGenlenti_N-MYC), were designed and cloned by GenScript (Piscataway, NJ). All constructs were generated by inserting the respective transgene fragments into the pGenlenti backbone. The respective lentiviruses were produced at the Cincinnati Children’s Hospital Viral Vector Core using the BRAF and CRAF constructs described above.

For lentiviral transduction, 5 × 10^4^ immortalized CAFs were seeded in one well of a 6-well plate. When the cells reached 50% confluence, 50 µL of virus was mixed with 450 µL of basal DMEM supplemented with 10 μg/mL polybrene (Santa Cruz, sc-134220). After a 16-hour incubation, the virus medium was replaced with 2 mL of fresh DMEM supplemented with 10% FBS for cell culture. Depending on the antibiotic resistance gene cloned in each lentiviral construct, transduced cells were selected using 10 µg/mL puromycin (Gibco, A11138--03), 50 µg/mL hygromycin B, or 10 μg/mL blasticidin (Santa Cruz Biotechnology, sc-495389). The cells were incubated with antibiotics for 7 days and then transferred to normal DMEM.

### siRNA transfection

Silencer® siRNAs targeting ARAF (ID: 153), BRAF (ID:507), CRAF (ID: 1548), NRAS (ID: 120250), KRAS (ID: 120703), HRAS (ID 120898), and scramble Silencer® siRNA control (Cat# 4390843) were purchased from Thermo Fisher Scientific (Rochester, NY). CAFs expressing BRAF-WT or BRAF(T529N) were transfected with FlexiTube siRNAs targeting BRAF (ID: 673) to silence endogenous BRAF expression. CAFs expressing CRAF-WT or CRAF(T421N) were transfected with FlexiTube siRNAs targeting CRAF (ID: 5894) to silence endogenous CRAF expression. FlexiTube siRNAs targeting the 3′ untranslated region (3′ UTR) of BRAF and CRAF were obtained from Qiagen (Hilden, Germany) Briefly, 1 × 10^5^ CAFs were seeded in one well of a 6-well tissue culture plate and allowed to grow to approximately 80% confluence. Prior to transfection, culture medium was replaced with fresh DMEM. The cells were then transfected with the indicated siRNAs using Lipofectamine RNAiMAX (Thermo Fisher, Rochester, NY). For each transfection, a siRNA-Lipofectamine mixture was prepared by incubating 9 μL of Lipofectamine RNAiMAX and 30 pmol of the indicated siRNA in 300 μL of DMEM at room temperature for 5 minutes. The cells in each well were then incubated with siRNA-Lipofectamine mixture for 48 hours at 37°C in a humidified incubator with 5% CO₂. Afterwards, the transfection mixture was then replaced with DMEM. To silence NRAS, KRAS, and HRAS expression in CAFs, three siRNAs targeting NRAS, KRAS, and HRAS (30 pmol each as described above) were mixed and incubated with 9 μL of Lipofectamine RNAiMAX in 300 μL DMEM for transfection.

### CAF treatment

To treat CAFs with small-molecule compounds, 5 × 10^3^ iM27 or iM50 cells were seeded in each well of an 8-well Nunc™ Lab-Tek™ II chamber slide (Thermo Fisher Scientific, Rochester, NY) and cultured for three days. The cells were treated as indicated: 2 μM PLX4032 for 72 hours; 0.5 μM GSK2118436 for 72 hours; 50 nM jasplakinolide for 2 hours; 500 nM cytochalasin D for 24 hours; 10 μM Y-27632 for 24 hours; and 10 μM HA-1077 for 24 hours or 0.5 μM SCH772984 for 24 hours. Following treatment, the cells were subjected to immunofluorescence staining directly on slides.

### Chamber slide immunofluorescence staining

After treatment, the cells in each chamber were fixed in 4% paraformaldehyde for 10 minutes at room temperature and permeabilized with 0.05% Triton X-100 for 10 minutes on ice for immunofluorescence staining. After permeabilization, the cells were washed three times with PBS and blocked with 10% BSA for one hour at room temperature. Primary antibodies for SUN2 (Millipore, HPA001209, 1:200), Nesprin-2 (Abcam, clone EPR28137-54, 1:200), lamin A/C (Proteintech, 10298-1-AP, 1:200), F-actin (Thermo Fisher, clone R415, 1:200), β-catenin (Thermo Fisher, clone 15B8, 1:500), paxillin (BD Biosciences, clone 349, 1:200), and α-SMA (Thermo Fisher, clone 1A4, 1:200) were then added to each chamber and incubated overnight at 4°C. The next day, after washing with PBS three times, an Alexa Fluor 488- or 594-conjugated secondary antibodies (Thermo Fisher, Rochester, NY) were added to the corresponding well for a one-hour incubation at room temperature. Slides were counterstained with Hoechst 33342 (Hoechst), mounted with VECTASHIELD Antifade Mounting Medium (Vector Laboratories, Burlingame, CA), and coverslipped. Images were acquired using a Nikon Eclipse 80i fluorescence microscope or Zeiss LSM 710 confocal microscope. To measure nuclear β-catenin intensity, the nuclei were stained with Hoechst at a concentration of 10 µg/mL for 10 minutes at room temperature. Nuclei were identified based on Hoechst fluorescence, and the β-catenin signal within these regions was quantified using the ImageJ software.

### Cell stiffness experiments

To evaluate how CAFs respond to stiff matrix environments, CAFs were cultured for 24 hours on soft slides with either 0.2 kPa or 50 kPa stiffness (Matrigen, Irvine, CA). Cell culture, treatment, and immunofluorescence staining were performed using the same protocols described above.

### Proximity ligation assay (PLA)

PLA was performed using the Duolink® *In Situ* Red Starter Kit (DUO92101, Millipore, Burlington, MA) according to the manufacturer’s instructions. Briefly, 0.5 × 10^4^ cells were cultured in each well of an 8-well Nunc™ Lab-Tek™ II chamber slide. After culturing for three days under the indicated conditions, the cells were fixed in 4% paraformaldehyde for 10 minutes at room temperature and permeabilized with 0.1% Triton X-100 in PBS for 10 minutes on ice. Afterwards, the cells were blocked with Duolink blocking solution for one hour at 37°C and incubated overnight at 4°C with the following primary antibodies: BRAF (Proteintech, 20899-1-AP), CRAF (Proteintech, clone 1D6A1), FLAG (Proteintech, 20543-1-AP), or MYC (Proteintech, clone 1A5A2). The next day, the slides were washed with wash buffer A and incubated with species-specific PLA probes (PLUS and MINUS) for one hour at 37°C. After probe incubation, the cells were washed twice with wash buffer A for 5 minutes each. Ligation-ligase solutions were then added to each well and incubated at 37°C for 30 minutes to ligate the two probes into a circular DNA template. The cells were subsequently washed twice with wash buffer A for two minutes each. The circular templates were amplified by adding polymerase in amplification buffer containing nucleotides and fluorescently labeled oligonucleotides, resulting in distinct fluorescent spots. The slides were then mounted with Duolink *in* situ mounting medium containing DAPI for 30 minutes and imaged using a Nikon Eclipse 80i fluorescence microscope. PLA signals were quantified using ImageJ as either the average dot intensity per 40X field or number of dots per cell.

### Coimmunoprecipitation (co-IP)

Co-IP was performed using a Pierce Crosslink Immunoprecipitation Kit according to the manufacturer’s instructions (Thermo Fisher, Waltham, MA). Briefly, 10 μg of Myc-tag antibody (Proteintech, clone 1A5A2) or mouse IgG (Thermo Fisher, 31903) was added to a spin column containing 20 μL of resin slurry in a 1.5 mL collection tube. The antibodies and beads were incubated at room temperature with gentle rotation for one hour. After incubation, the spin columns were centrifuged, and the antibody-bound resin was subsequently rinsed with 1X coupling buffer three times to remove unbound antibodies. A 2.5 mM disuccinimidyl suberate (DSS) crosslinker solution was added to crosslink the antibody to the resin. Subsequently, 1 mg of protein extracted from the cells co-expressing Flag-tagged CRAF and Myc-tagged CRAF with or without PLX4032 treatment was added to a spin column and incubated with rotation at 4°C overnight. After incubation, the column was washed three times with 1X TBS and one time with 1X conditioning buffer. The samples were eluted in 50 μL of elution buffer. The eluate was boiled with sample buffer at 95°C for five minutes for Western blotting.

### Western blotting

For Western blotting, 20 × 10⁴ iM27 or iM50 cells were seeded in one 10 cm culture dish and treated as described above for three days. After treatment, proteins were extracted using RIPA buffer supplemented with EDTA-free Halt Protease and Phosphatase Inhibitor Cocktail (Thermo Fisher, 78443). Briefly, cells were lysed in RIPA buffer on ice for 30 minutes with brief vortexing every 10 minutes. The lysates were then centrifuged at 15,000 × g for 15 minutes at 4°C, and the supernatants were collected and transferred to fresh tubes. Protein was quantified using the Bicinchoninic Acid (BCA) protein assay (Thermo Fisher, A55860). Twenty μg of protein were loaded per lane and run on a 10-12% SDS‒PAGE gel and subsequently transferred to a nitrocellulose membrane. The membranes were blocked with 5% fat-free milk or 5% BSA in TBST for one hour at room temperature and then incubated with the following primary antibodies: ARAF (Cell Signaling Technology, 4432, 1:1000), BRAF (Proteintech, 20899-1-AP, 1:1000), CRAF (Proteintech, clone 1D6A1, 1:1000), CRAF (Proteintech, 26863-1-AP, 1:1000), phospho-BRAF (Cell Signaling Technology, 2696, 1:1000), phospho-CRAF (Cell Signaling Technology, 9127, 1:1000), Flag (Proteintech, 20543-1-AP, 1:1000), MYC (Proteintech, clone 1A5A2, 1:1000), MYC (Cell Signaling Technology, clone 71D10, 1:1000), KRAS (Proteintech, 12063-1-AP, 1:1000), HRAS (Proteintech, 18295-1-AP, 1:1000), NRAS (Proteintech, 10724-1-AP, 1:1000), ERK1/2 (Cell Signaling Technology, clone 3A7, 1:1000), and phospho-ERK (Cell Signaling Technology, clone D13.14) in blocking buffer. After washing three times with TBST, the membranes were then incubated with IRDye 800CW donkey anti-rabbit IgG secondary antibody (LI-COR, 926–32213, 1:2000) or IRDye 680RD donkey anti-mouse IgG secondary antibody (LI-COR, 926–68072, 1:2000) at a 1:2000 dilution in blocking buffer and washed three times with TBST. The membranes were imaged using an Odyssey CLx imaging system (LI-COR, model #9140).

### Quantification of nuclear morphology

First, 0.5 × 10⁴ CAFs were seeded in one well of an 8-well chamber slide and treated as indicated in each figure. Following treatment, nuclei were stained with Hoechst, and confocal images of randomly selected fields were captured at 40X magnification using a Zeiss LSM 710 confocal microscope. Hoechst-stained images were analyzed using ImageJ software. The aspect ratio and circularity of individual nuclei were quantified using the Analyze Particles function.

### Actin cap quantification

A total of 0.5 × 10⁴ CAFs seeded in one well of an 8-well chamber slide were stained for F-actin using Phalloidin Labeling Probes (Thermo Fisher, R415, 1:200), and nuclei were stained with Hoechst. Confocal images of randomly selected fields were acquired at 40X magnification using a Zeiss LSM 710 confocal microscope. Actin caps were identified as dense F-actin fibers spanning over the apical surface of the nucleus. The intensity of F-actin staining of the actin caps was quantified using the ImageJ software.

### Z-stack confocal microscopy

To visualize the three-dimensional organization of actin caps relative to the nucleus, Z-stack images were acquired at 40X magnification using a Zeiss LSM 710 confocal microscope. Hoechst staining was used to define nuclear boundaries. A single cell was scanned from the bottom to the top of the nucleus at 1 µm intervals, covering the entire nuclear volume.

### Quantification and statistical analysis

Data were analyzed using the GraphPad Prism 9 software package (GraphPad Software Inc., San Diego, CA) and expressed as the mean ± SD. The mean difference was determined using Student’s t-test and considered statistically significant at P < 0.05.

## RESULTS

### BRAFis induce nuclear deformation in CAFs

CAFs translate therapeutic and mechanical pressures into transcriptional reprogramming, which fuels tumor progression and underpins resistance to therapy. However, the precise mechanisms by which therapeutic agents drive CAF activation and crosstalk with mechanical signaling pathways remain to be fully elucidated. Nuclear morphological changes are known mechanism by which cells regulate gene expression^29^. To understand how CAFs adapt to BRAFis, we first examined the nuclear shape of CAFs after treatment with PLX4032 (Fig. 1) and GSK2118436 (Fig. S1). As shown in Figs. 1A and S1A, the nuclei in many BRAFi-treated CAFs became elongated. The average nuclear aspect ratio in BRAFi-treated CAFs, which was calculated by dividing the length of the longest axis of the nucleus by the length of its shortest axis, was ∼1.94, compared with ∼1.39 in DMSO-treated CAFs (Figs. 1B and S1B), suggesting that BRAFis stretched the nuclei in CAFs. Furthermore, nuclear circularity, which represents the roundness of a nucleus, was reduced in BRAFi-treated CAFs to ∼0.63 from ∼0.77 in DMSO-treated CAFs (Figs. 1C and S1C). These observations indicate that BRAFis caused the nuclei in CAFs to lose their typical shape.

**Figure 1.**
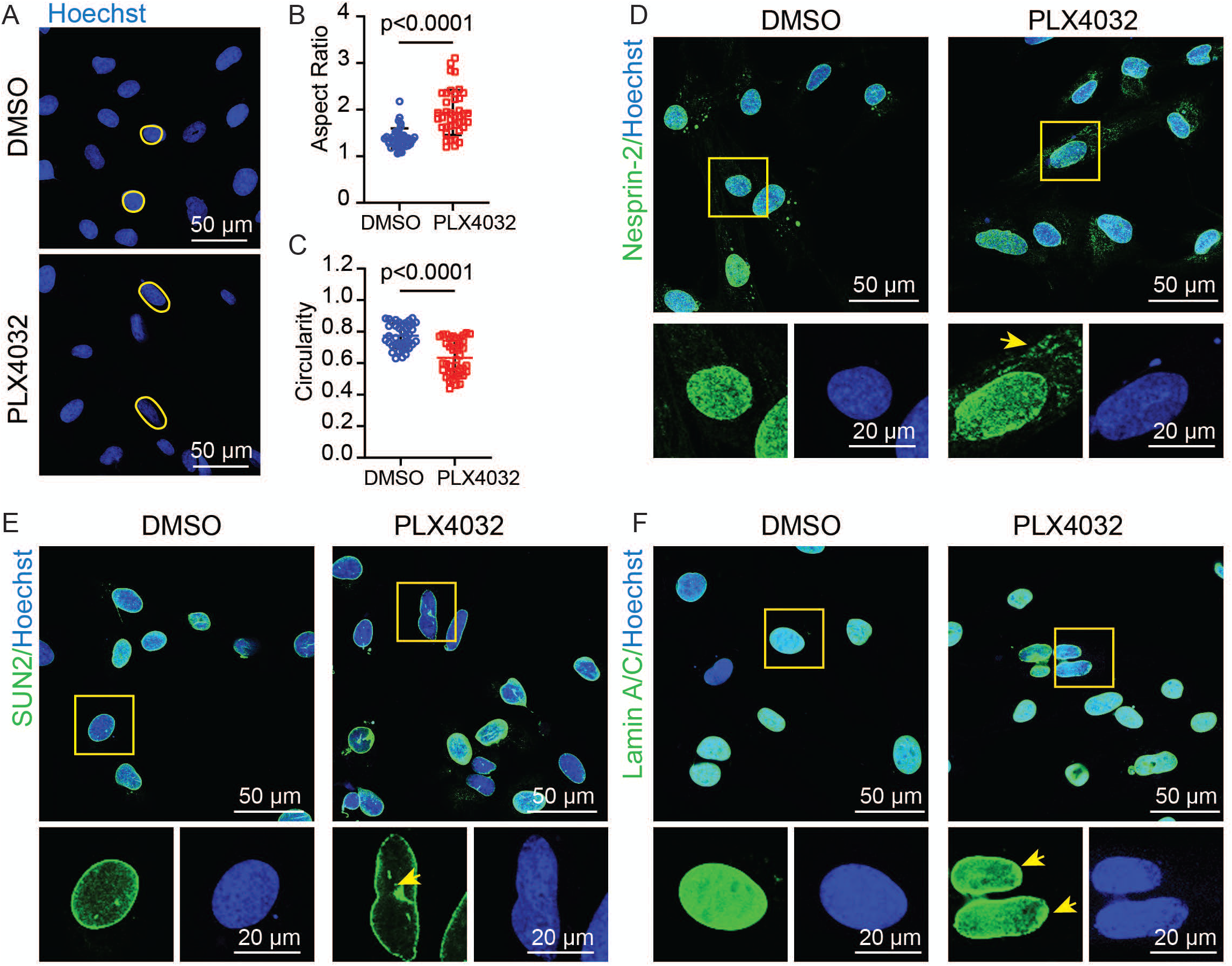
BRAFis induce nuclear deformation in CAFs. (A) Representative confocal images showing nuclei stained with Hoechst in iM27 with or without PLX4032 treatment. Yellow circles highlight the approximate nuclear boundaries. iM27 cells were treated with either DMSO or PLX4032 for comparison. (B–C) Scatter dot plots showing the nuclear morphological parameters in iM27 with or without PLX4032 treatment. (B) The nuclear aspect ratio and (C) circularity were analyzed and quantified from confocal images using ImageJ. The data are presented as mean ± SD (n = 40-46 nuclei per group). (D–F) Confocal images showing immunostaining of key components of the LINC complex in iM27 w/ or w/o PLX4032 treatment, including (D) Nesprin-2, (E) SUN2, and (F) nuclear lamin A/C. In each figure, individual cells shown in small boxes are representative stained cells, denoted by yellow boxes in the large boxes above. Disrupted distribution patterns of Nesprin-2, SUN2, and lamin A/C in PLX4032-treated iM27 cells are indicated by yellow arrows.

The nuclear shape and positioning are regulated by the linker of the nucleoskeleton and cytoskeleton (LINC) complex, which connects the nucleus (nucleoskeleton) to the cytoskeleton^30^. Alterations in the LINC complex in BRAFi-treated CAFs were visualized by immunofluorescence staining of two core LINC components, Nesprin-2 and SUN2. Nesprin-2 is anchored in the outer nuclear membrane and interacts with actin filaments, thereby joining the cytoskeleton to the nucleoskeleton^31^. As shown in Figs. 1D and S1D, Nesprin-2 in CAFs extend into the cytoplasm. This structural arrangement enables the LINC complex to sense and transmit mechanical and associated signaling cues from the cytoplasm to the nucleus. Interestingly, upon BRAFi treatment, cytoplasmic localization of Nesprin-2 surrounding the nucleus was increased in CAFs. SUN2 is a membrane-embedded protein located in the inner nuclear membrane, where it interacts with nuclear lamins such as lamin A/C^32^. BRAFi-treated CAFs presented aberrant nuclear distribution of SUN2, suggesting disruption of the LINC complex and possible compromise of the nuclear envelope (Figs. 1E and S1E). Since the nuclear membrane was disrupted, we next investigated whether there were changes in nuclear lamins, as they are directly connected to the inner nuclear membrane and are essential structural proteins^33^. Discontinuous staining of lamin A/C in BRAFi-treated CAFs confirmed the disruption of the nuclear lamin structure (Figs. 1F and S1F). The findings suggest that changes in the LINC complex may contribute to the nuclear deformation in BRAFi-treated CAFs.

### BRAFis trigger actin polymerization and mechanically sculpt nuclear architecture

Actin polymerization, particularly around the nucleus, plays an important role in regulating the LINC complex and the nuclear shape^34^. It is possible that the increased actin polymerization exerts a mechanical force on the nucleus and can cause nuclear rupture. To understand whether the BRAFi-induced changes in the LINC complex and nuclear elongation observed in CAFs were caused by actin-mediated nuclear force, we investigated actin filament organization in PLX4032-treated CAFs. As shown in Fig. S2A, B, the number of polymerized actin filaments was greater in CAFs than in normal dermal fibroblasts. However, the actin structures were much more prominent in CAFs treated with PLX4032 (Fig. S2C, D), suggesting that BRAFis induced cytoskeletal remodeling in CAFs. We focused on the perinuclear actin cap, a specialized subset of actin filaments that span the nucleus^35^, because it is directly anchored to the LINC complexes on the nuclear envelope. Z-stack confocal imaging revealed prominent assembly of actin cap structures in PLX4032-treated CAFs, where F-actin aligned above the nucleus in a dome-like structure and colocalized with nuclear staining, as indicated by the yellow arrow (Fig. S2E).

These observations suggest that mechanical forces exerted by enhanced actin cytoskeleton remodeling, which is induced by BRAFis, may contribute to nuclear deformation in CAFs. To test this hypothesis, we treated CAFs with jasplakinolide (jaspla), an actin polymerization inducer known to promote cytoskeletal remodeling^36^. As shown in Fig. 2A, B, actin polymerization and the formation of perinuclear actin caps were significantly increased in jaspla-treated CAFs. Interestingly, Jaspla treatment also induced nuclear deformation (Fig. 2C). Consistent with PLX4032 treatment, Jaspla-treated CAFs showed an increase in the nuclear aspect ratio from ∼1.30 to 1.69 and a reduction in circularity from ∼0.83 to 0.68, indicating that the nuclei became elongated and less round (Fig. 2D). These data imply that increased cytoskeletal remodeling is directly responsible for the nuclear deformation in CAFs.

**Figure 2.**
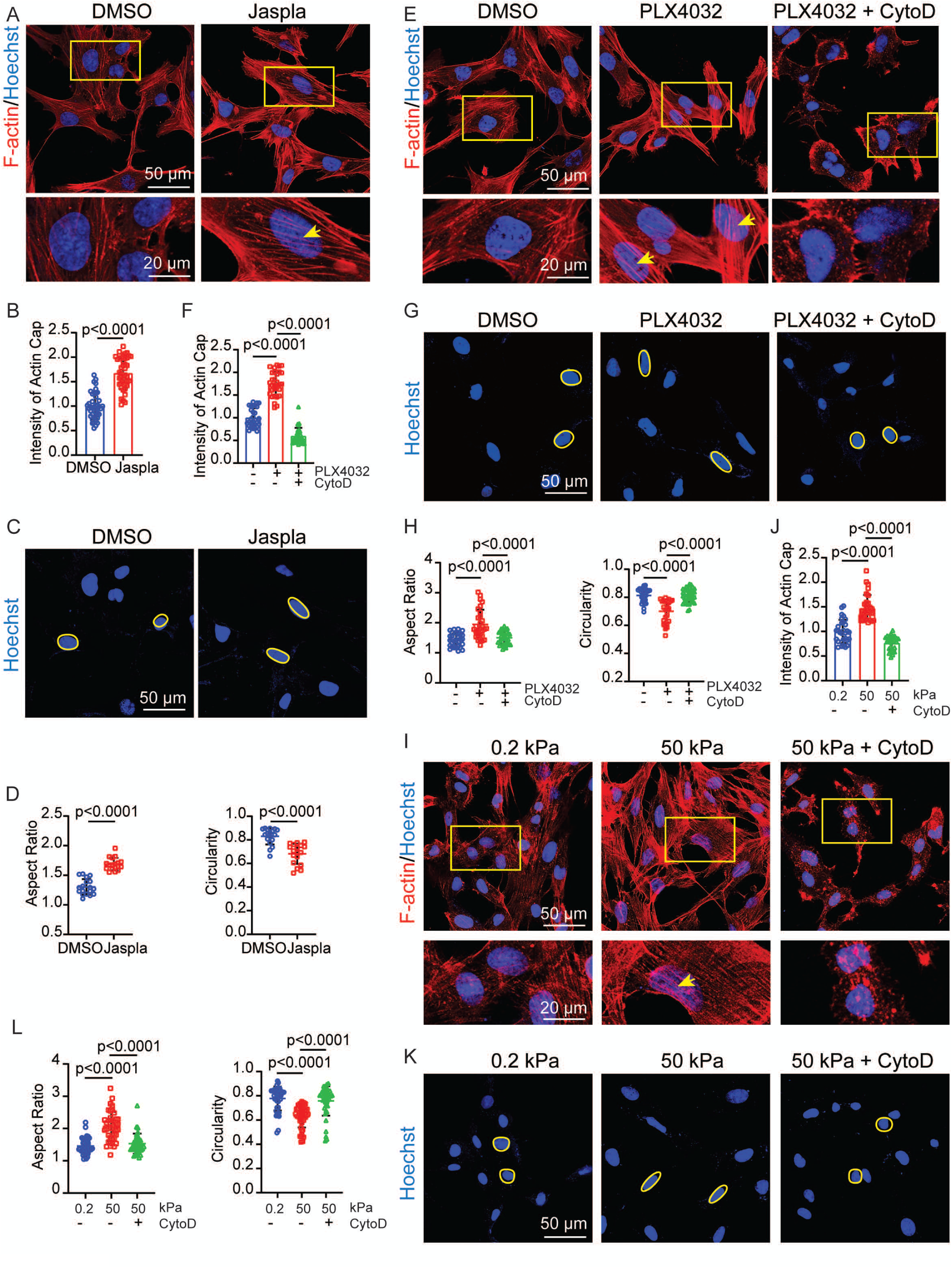
BRAFis and matrix stiffness trigger actin polymerization leading to nuclear deformation. (A) Confocal images of F-actin in iM27 cells treated with DMSO or Jaspla. The representative individual cells shown in the insets correspond to the cells marked by yellow boxes in the larger images above. The actin caps are indicated by yellow arrows. (B) Quantification of the actin cap intensity using ImageJ. The data are presented as mean ± SD (n = 42 nuclei per group). (C) Representative confocal images showing nuclei stained with Hoechst in iM27 cells treated with DMSO or Jaspla. Yellow circles highlight the approximate nuclear boundaries. (D) Quantification of the nuclear morphological changes, including the nuclear aspect ratio (left) and circularity (right), analyzed and quantified from confocal images using ImageJ. The data are presented as mean ± SD (n = 18 nuclei per group). (E) Confocal images of F-actin in iM27 cells treated with DMSO, PLX4032, or the combination of PLX4032 and CytoD. The representative individual cells shown in the insets correspond to the cells marked by yellow boxes in the larger images above. The actin caps are indicated by yellow arrows. (F) Quantification of the actin cap intensity using ImageJ. The data are presented as mean ± SD (n = 30 nuclei per group). (G) Representative confocal images of Hoechst-stained nuclei in iM27 cells treated with DMSO, PLX4032, or the combination of PLX4032 and CytoD. Yellow circles highlight the approximate nuclear boundaries. (H) Quantification of nuclear morphology, including the nuclear aspect ratio (left) and circularity (right), was performed, and the results were analyzed and quantified from confocal images via ImageJ. The data are presented as mean ± SD (n = 26-39 nuclei per group). (I) Confocal images of F-actin in iM27 cells cultured on soft slides with a stiffness of 0.2 kPa (left), hard slides with a stiffness of 50 kPa (middle), and hard slides with a stiffness of 50 kPa and CytoD treatment (right). The representative individual cells shown in the insets correspond to the cells marked by yellow boxes in the larger images above. The actin caps are indicated by yellow arrows. (J) Quantification of the actin cap intensity using ImageJ. The data are presented as mean ± SD (n = 25-30 nuclei per group). (K) Representative confocal images showing nuclei stained with Hoechst in iM27 cells cultured on soft slides with a stiffness of 0.2 kPa (left), hard slides with a stiffness of 50 kPa (middle), and hard slides with a stiffness of 50 kPa and CytoD treatment (right). Yellow circles highlight the approximate nuclear boundaries. (L) Scatter dot plots showing the nuclear morphological changes in iM27 among three different conditions. the nuclear aspect ratio (left) and circularity (right) were analyzed and quantified from confocal images using ImageJ. The data are presented as mean ± SD (n = 40 nuclei per group).

We also investigated whether inhibiting actin polymerization could reverse the changes in nuclear shape induced by PLX4032. To this end, CAFs were treated with PLX4032 alone or in combination with cytochalasin D (CytoD), an actin polymerization inhibitor^37^. F-actin staining confirmed a reduction in actin fiber polymerization and actin cap structure formation in CAFs treated with PLX4032 + CytoD compared to those treated with PLX4032 only (Fig. 2E, F), suggesting that CytoD effectively disrupted actin filament assembly driven by PLX4032. As predicted, PLX4032-induced nuclear deformation was significantly reversed by CytoD, with the nuclear aspect ratio decreasing to approximately 1.50 and the nuclear circularity increasing to approximately 0.81 (Fig. 2G, H).

### Matrix stiffness drives cytoskeletal polymerization and deforms the nucleus

Given that increased tissue stiffness is a classic characteristic of solid tumors, we tested whether elevated matrix stiffness promotes actin polymerization and nuclear deformation in CAFs. CAFs were cultured on either a soft substrate with a stiffness of 0.2 kPa or a hard substrate with a stiffness of 50 kPa. In line with the effects of PLX4032 stimulation, increased ECM stiffness induced marked increases in actin polymerization and actin cap formation in CAFs cultured on stiff substrates compared to soft substrates (Fig. 2I, J). This stiffness-induced cytoskeletal remodeling was abrogated by CytoD, mirroring its effect upon PLX4032 stimulation (Fig. 2E). Increased ECM stiffness also elevated the nuclear aspect ratio and reduced the nuclear circularity, suggesting that the nuclei were deformed. CytoD reversed these stiffness-driven nuclear changes, reducing the aspect ratio to ∼1.53 and restoring circularity to ∼0.76 (Fig. 2K, L). These findings suggest that ECM stiffness also elicits nuclear deformation by driving cytoskeleton remodeling, indicating a common force transmission mechanism in CAFs that is shared with BRAF inhibition.

### Nuclear deformation facilitates β-catenin entry into the nucleus

Next, we aim to understand how nuclear dynamics translates into molecular signals in CAFs. Previously, we reported that BRAFis induce Wnt-independent activation of nuclear β-catenin signaling in CAFs^7^. β-catenin is a dual-function protein that functions in cell adhesion at the cellular membrane and mediates signaling by shuttling between the cytoplasm and nucleus. Confocal imaging of SUN2-stained CAFs revealed that the nuclear distribution of SUN2 in jaspla-treated CAFs was similar to that observed in PLX4032-treated CAFs, with increased nuclear β-catenin accumulation, reflecting a positive correlation between nuclear deformation and β-catenin nuclear entry (Fig. 3A, C). In contrast, abnormal SUN2 localization in PLX4032-treated CAFs was diminished by simultaneous CytoD treatment (Fig. 3D). Consequently, CytoD also suppressed nuclear β-catenin accumulation in PLX4032-treated CAFs (Fig. 3E, F).

**Figure 3.**
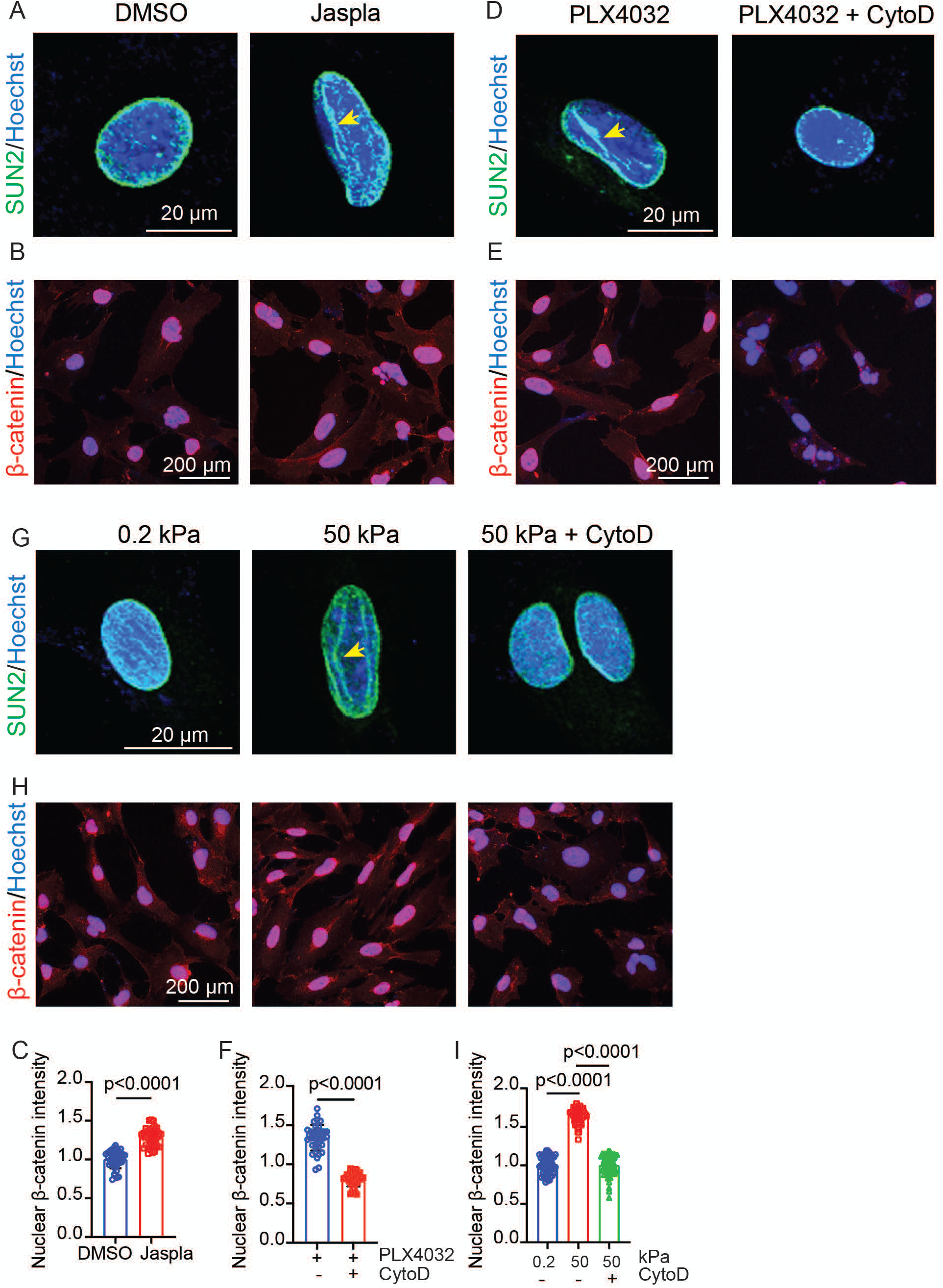
Nuclear deformation facilitates WNT-independent β-catenin entry into the nucleus. (A–B) Confocal images showing SUN2 staining (A) and fluorescence microscopy images showing nuclear β-catenin staining (B) in iM27 cells treated with DMSO or jaspla. Disrupted SUN2 distribution in jaspla-treated iM27 cells is indicated by yellow arrows. (C) Quantification of the nuclear β-catenin intensity in iM27 cells treated with DMSO or Jaspla. Data are presented as mean ± SD (n = 40 cells per group). (D–E) Confocal images showing SUN2 staining (D) and fluorescence microscopy images showing nuclear β-catenin staining (E) in iM27 cells treated with PLX4032 or the combination of PLX4032 and CytoD. Yellow arrows indicate the disorganized SUN2 distribution in iM27 cells treated with PLX4032 alone. (F) Quantification of the nuclear β-catenin intensity in iM27 cells treated with PLX4032 or the combination of PLX4032 and CytoD. Data are presented as mean ± SD (n = 40 cells per group). (G) Confocal images showing SUN2 staining in iM27 cells cultured on soft slides with a stiffness of 0.2 kPa (left), hard slides with a stiffness of 50 kPa (middle), and hard slides with a stiffness of 50 kPa and CytoD treatment (right). Disorganized SUN2 distribution in iM27 cells cultured on 50 kPa slides is indicated by yellow arrows. (H) Fluorescence microscopy images showing nuclear β-catenin staining in iM27 cells cultured on soft slides with a stiffness of 0.2 kPa (left), hard slides with a stiffness of 50 kPa (middle), and hard slides with a stiffness of 50 kPa and CytoD treatment (right). (I) Quantification of the nuclear β-catenin intensity in iM27 cells under three different conditions. Data are presented as mean ± SD (n = 60 cells per group).

Notably, the abnormal nuclear localization of SUN2 observed on stiff substrates was also rescued by CytoD treatment (Fig. 3G). In parallel, the increased nuclear accumulation of β-catenin in CAFs cultured on the hard substrate compared with the soft substrate was markedly reduced upon CytoD treatment (Fig. 3H, I). Collectively, these data demonstrate that nuclear deformation acts as a key facilitator of β-catenin nuclear transport driven by both increased matrix stiffness and BRAFis.

### Nuclear reshaping drives an actin-β-catenin feedback loop to reinforce CAF activation

Nuclear β-catenin is crucial for regulating the expression of genes that are essential for cell proliferation, differentiation, and survival^38^. Since β-catenin is shuttled into the nucleus by various stimuli, including BRAFis and stiff substrates, we wanted to understand the biological outcome of increased nuclear β-catenin in fibroblasts and melanoma development. As shown in Fig. S3A and B, fibroblasts overexpressing a constitutively active form of β-catenin (*Co1α2-CreER^T2^; Rosa-rtTA; TetO-ΔN-β-catenin*, hereafter ΔN-β-catenin) displayed higher nuclear β-catenin expression than control fibroblasts expressing endogenous β-catenin (*Co1α2-CreER^T2^; Rosa-rtTA*, hereafter control). These ΔN-β-catenin-overexpressing fibroblasts exhibited increased proliferation (Fig. S3C) and elevated expression of α-SMA, which is a well-established CAF marker (Fig. S3D, E). Furthermore, the expression and distribution of paxillin and F-actin were upregulated in ΔN-β-catenin-expressing fibroblasts (Fig. S3F-I). Since changes in the actin cytoskeleton and scaffolding proteins such as paxillin are often linked to cell contractility, we assessed their contractile capacity using a gel contraction assay and confocal reflection microscopy (CRM) (Fig. S3J-N). The results showed that ΔN-β-catenin-expressing CAFs exhibited increased contractility. Importantly, as noted above, increased actin polymerization further amplifies nuclear deformation and β-catenin accumulation, establishing a reciprocal feed-forward loop that can continuously enhance the biological properties of fibroblasts.

To investigate the importance of ΔN-β-catenin-expressing CAFs in *in vivo* melanoma progression, we co-injected fibroblasts and BRAF-mutant D4M melanoma cells intradermally into B6 mice to induce melanoma formation (Fig. 4A). Melanomas containing ΔN-β-catenin-expressing CAFs (ΔN-β-catenin melanomas) grew faster than those containing wild-type CAFs at each time point (Fig. 4B). On day 18, when tumors were collected, melanomas containing ΔN-β-catenin-expressing CAFs were larger and weighed more than control melanomas (Fig. 4C, D), suggesting that upregulated β-catenin activity in CAFs promoted melanoma growth *in vivo*. H&E staining revealed that the structure of the ΔN-β-catenin melanomas was more compact with fewer intercellular spaces than that of the control tumors (Fig. 4E). To confirm this observation, we assessed the expression of two major proliferation markers, Ki67 and Cyclin D1. As shown in Fig. 4F and G, the number of Ki67+ melanoma cells (α-SMA-) was notably increased in ΔN-β-catenin melanomas, suggesting that upregulated β-catenin activity in CAFs promoted the proliferative activity of BRAF-mutant melanoma cells. In line with this observation, increased expression of Cyclin D1 was detected in ΔN-β-catenin melanomas (Fig. 4H, I).

Increased α-SMA expression was observed in these melanomas, indicating that upregulated β-catenin activity drove CAF activation (Fig. 4J, K), which was consistent with the *in vitro* data showing that β-catenin overexpression promoted fibroblast proliferation (Fig. S3C). CAFs are the major producers of ECM components, such as fibronectin and collagen, which play critical roles in remodeling the TME and promoting tumor malignancy. Increased fibronectin and collagen deposition were observed in ΔN-β-catenin melanomas (Fig. 4L-O). In conclusion, increased nuclear β-catenin, which can be driven by BRAFis or stiff stroma, reprograms CAFs and sustains their activation, playing a critical role in promoting matrix remodeling and melanoma progression.

### BRAFi binds to the RAF kinase domain and promotes BRAF and CRAF dimerization and phosphorylation

We then investigated how BRAFi treatment reorganizes the cytoskeleton to promote nuclear deformation and β-catenin entry in CAFs. The RAF kinase family includes three serine/threonine-specific protein kinases, ARAF, BRAF, and CRAF^39^. We used small interfering RNA (siRNA) to selectively silence each RAF kinase to determine which RAF isoforms mediate the response to PLX4032 in CAFs (Fig. 5A). As shown in Fig. 5B and C, ARAF depletion had no significant effect on PLX4032-induced nuclear accumulation of β-catenin compared with that in mock-infected controls. In contrast, loss of either BRAF or CRAF markedly suppressed β-catenin nuclear entry in response to PLX4032. These findings were confirmed in the additional CAF lines (Fig. S4A–C). Together, these observations confirmed that BRAF and CRAF, but not ARAF, are involved in mediating the nuclear accumulation of β-catenin driven by BRAFis in CAFs.

**Figure 4.**
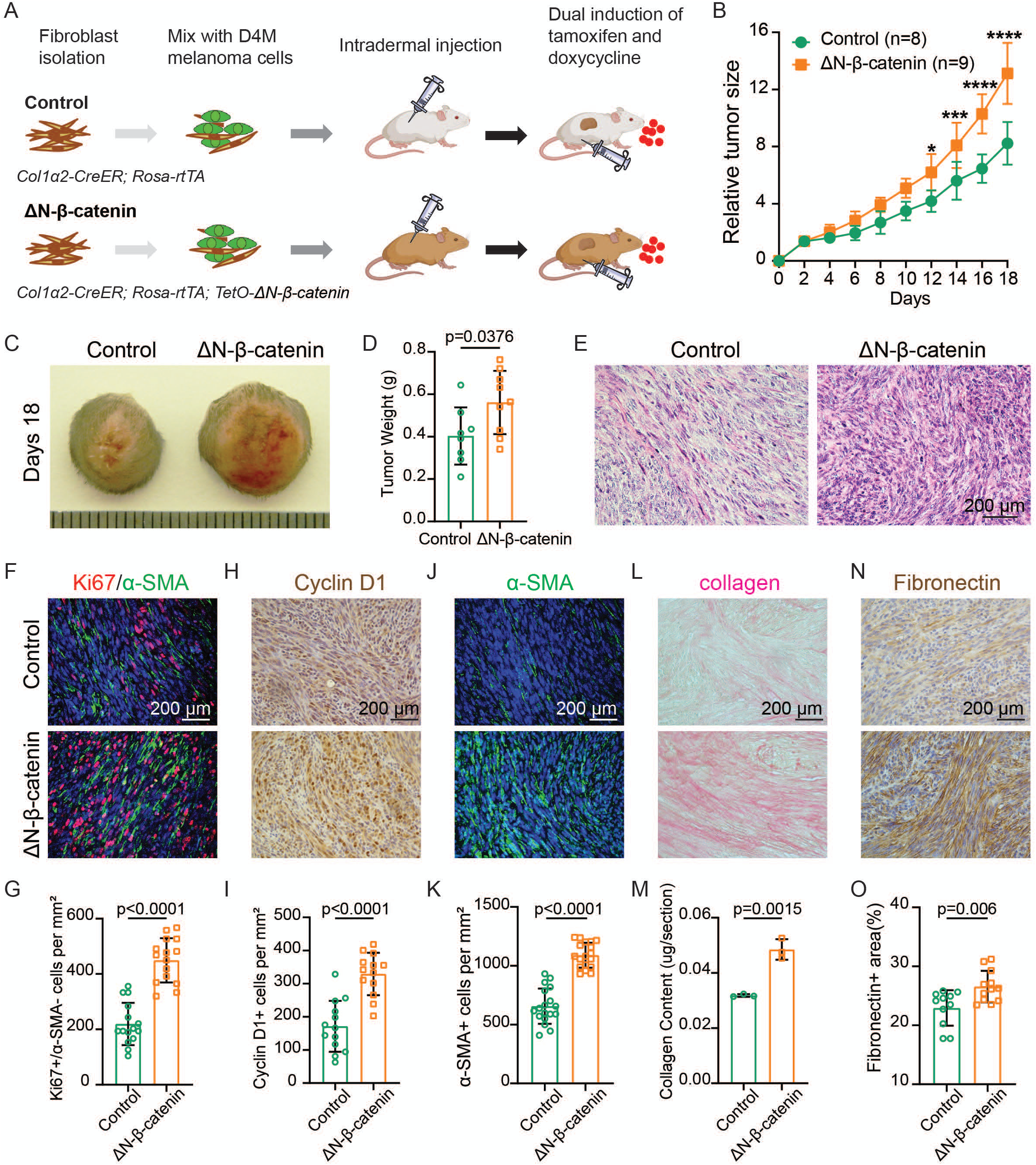
Nuclear deformation drives an actinβ-catenin feedback loop to reinforce CAF activation. (A) Schematic illustration of the melanoma mouse model used to study the *in vivo* effects of β-catenin-overexpressing fibroblasts on melanoma progression. D4M melanoma cells (green) were mixed with uninduced control fibroblasts (*Col1α2-CreER; Rosa-rtTA*, control melanoma) or mutant fibroblasts (*Col1α2-CreER; Rosa-rtTA; TetO-ΔN-β-catenin*, ΔN-β-catenin melanoma) and injected intradermally into the flanks of B6 mice. The tumors were allowed to grow until they reached approximately 62.5 mm³ in volume. At this point (designated day zero), tamoxifen and doxycycline were administered to induce β-catenin overexpression in the mutant fibroblasts. Tumor growth was subsequently monitored every other day until the end point. (B) Tumor sizes were measured every other day and compared between control melanomas and ΔN-β-catenin melanomas (n = 8-9). (C) Representative images of control melanomas and ΔN-β-catenin melanomas on day 18. (D) Comparison of tumor weight between control melanomas and ΔN-β-catenin melanomas on day 18. Each data point represents the weight of an individual tumor. The data are presented as the mean ± SD (n = 8-9). (E) H&E-stained control and ΔN-β-catenin melanoma tissue sections captured under a light microscope. (F). Images show α-SMA and Ki67 immunofluorescence staining of control and ΔN-β-catenin melanoma tissue sections as indicated. Nuclei were counterstained with DAPI (blue). (G) Comparison of the numbers of α-SMA-cells that were Ki67+ in each group. Each data point represents the number of α-SMA-; Ki67+ cells per mm^2^ were counted for each melanoma sample. n = 15. (H) Representative images of cyclin D1 staining of control and ΔN-β-catenin melanoma tissue sections from the indicated groups by immunohistochemistry. (I) The graph shows the numbers of cyclin D1+ cells in each group. Each data point represents the number of cyclin D1+ cells per mm^2^ counted for each melanoma sample. n = 13. (J) Images showing α-SMA staining of control and ΔN-β-catenin melanoma tissue sections from respective group. The nuclei were counterstained with DAPI (blue). (K) The graph shows the number of α-SMA+ cells in each group. Each data point represents the number of α-SMA cells per mm^2^ counted for each melanoma sample. n = 18. (L) Images showing collagen staining of melanoma tissue sections from the indicated groups. (M) Quantitative comparison of collagen contents in control and ΔN-β-catenin melanoma tissue sections by collagen extraction and colorimetric measurement. n = 3. (N) Immunohistochemistry images showing the expression of the ECM protein fibronectin in control and ΔN-β-catenin melanoma tissue sections as indicated. (O) Quantitative comparison of the percentages of the surface areas occupied by fibronectin+ fibroblasts measured in control and ΔN-β-catenin melanoma tissue sections. Eleven to twelve random fields from three melanoma pairs were counted. n = 11-12. For all the staining images, the scale bar represents 200 µm. In all the statistical graphs, the data are presented as mean ± SD. *, P<0.05; **, P<0.01; ***, P<0.001; ns, not significant.

**Figure 5.**
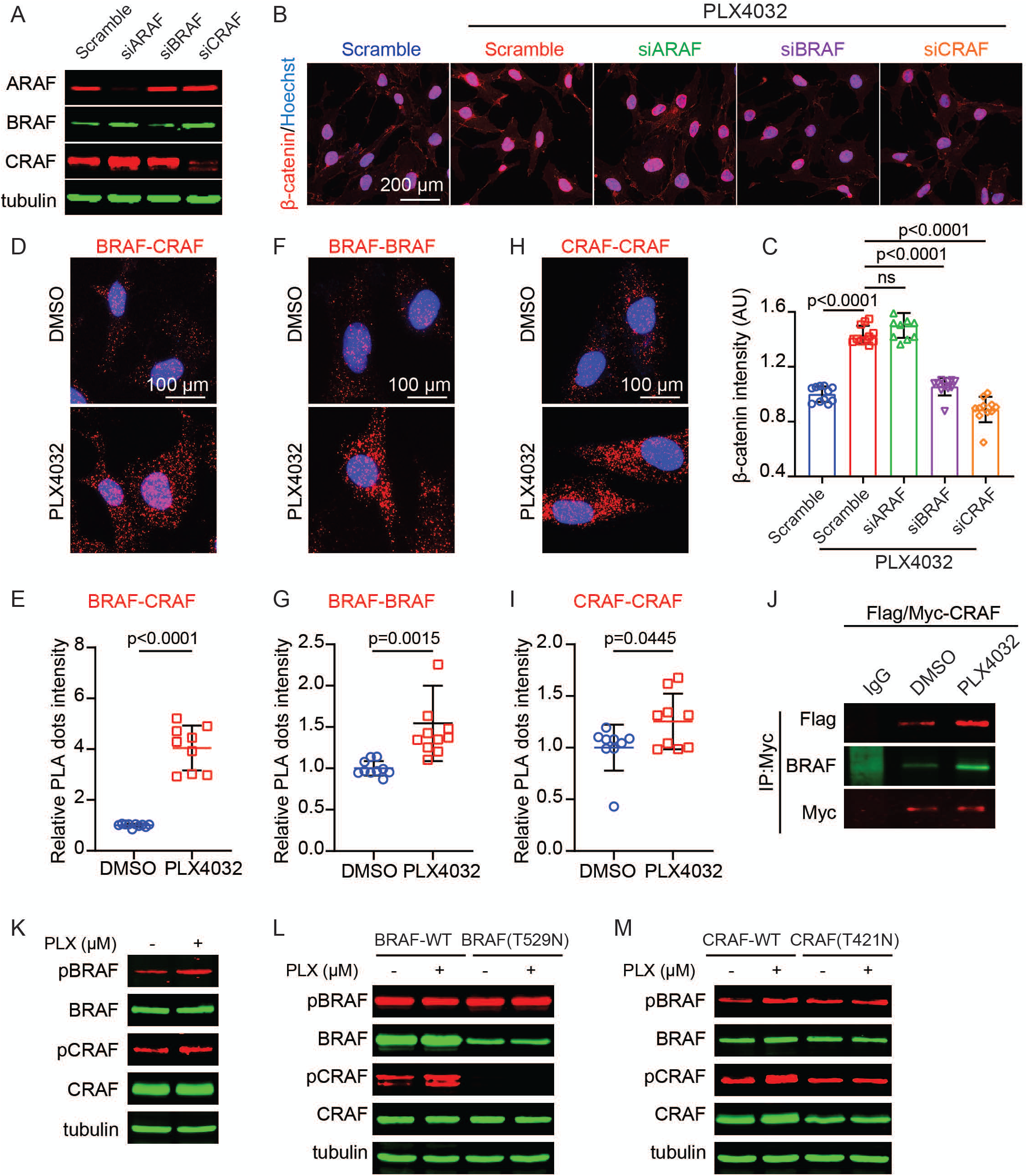
BRAFis bind to the RAF kinase domain and promote BRAF and CRAF dimerization and phosphorylation. (A) Western blot confirming effective silencing of ARAF, BRAF, or CRAF expression in iM27 cells using corresponding siRNAs. (B) Representative fluorescence microscopy images showing β-catenin expression in iM27 cells transfected with scramble siRNA or siRNAs that deplete ARAF, BRAF, or CRAF expression under PLX4032 treatment. iM27 transfected with scramble siRNA without PLX4032 treatment were used as a control. (C) Quantification of nuclear β-catenin intensity in iM27 and genetically modified iM27 cells as shown in (B) with or without PLX4032 treatment using ImageJ. The data are presented as mean ± SD (n = 11 randomly selected 20 X fields per group). (D) Representative PLA images showing BRAF-CRAF heterodimerization in DMSO- and PLX4032-treated iM27 cells. Red dots indicate BRAF-CRAF dimers. (E) Quantification of PLA signals comparing PLX4032-treated iM27 cells with DMSO-treated iM27 cells. The data are presented as mean ± SD (n = 9-10 randomly selected 40X fields per group). (F) Representative PLA images showing BRAF-BRAF homodimerization in DMSO- and PLX4032-treated iM27 cells. The red dots indicate BRAF-BRAF dimers. (G) Quantification of PLA signals comparing PLX4032-treated iM27 cells with DMSO-treated iM27 cells. The data are presented as mean ± SD (n = 9-10 randomly selected 40X fields per group). (H) Representative PLA images showing CRAF-CRAF homodimerization in DMSO- and PLX4032-treated iM27 cells. The red dots indicate CRAF-CRAF dimers. (I) Quantification of PLA signals comparing PLX4032-treated iM27 cells with DMSO-treated iM27 cells. The data are presented as the mean ± SD (n = 9-10 randomly selected 40X fields per group). (J) Western blot analysis showing increased CRAF-CRAF and CRAF-BRAF interactions in PLX4032-treated iM27 cells compared with those in DMSO-treated iM27 cells. Co-IP was performed via the use of an anti-Myc antibody to pull down interacting proteins in iM27 cells coexpressing Myc-tagged and Flag-tagged CRAF. (K–M) Western blot analysis of phosphorylated BRAF (Ser445) and CRAF (Ser338) expression in iM27 cells (K), BRAF-deficient iM27 cells overexpressing wild-type BRAF or BRAF (T529N) (L), and CRAF-deficient iM27 cells overexpressing wild-type CRAF or CRAF (T421N) (M) treated with either DMSO or PLX4032.

The activation of RAF isoforms is intrinsically linked to their ability to form dimers, either homodimers (e.g., BRAF-BRAF or CRAF-CRAF) or heterodimers (e.g., BRAF-CRAF), depending on the cellular context. To understand how BRAFi induces paradoxical RAF activation in CAFs, we first examined RAF dimerization using proximity ligation assays (PLA). PLA revealed greater BRAF-CRAF heterodimer formation in PLX4032-treated CAFs than in control cells (Figs. 5D, E and S4D, E). To investigate RAF homodimer formation in CAFs, we co-expressed Flag-tagged BRAF and Myc-tagged BRAF to identify BRAF-BRAF homodimerization, and in a separate set of cells, co-expressed Flag-tagged CRAF and Myc-tagged CRAF to assess CRAF-CRAF homodimerization. PLX4032 treatment notably enhanced the formation of both BRAF-BRAF and CRAF-CRAF homodimers (Fig. 5F-I). Additionally, the formation of BRAF and CRAF dimers was confirmed using another BRAFi, GSK2118436 (Fig. S4F-I). Coimmunoprecipitation (co-IP) using an anti-Myc antibody further confirmed increased CRAF homodimerization and heterodimerization in PLX4032-treated CAFs co-expressing Flag- and Myc-tagged CRAF (Fig. 5J). Furthermore, phosphorylation of BRAF at Ser445 and CRAF at Ser338, established markers of RAF activation, was elevated in CAFs following PLX4032 treatment (Fig. 5K). Together, these findings suggest that BRAFis promote RAF activation in CAFs by increasing RAF dimerization and phosphorylation.

In melanoma cells harboring the BRAF V600E mutation, PLX4032 binds to the kinase domain of the mutant BRAF to block its enzymatic activity. However, whether PLX4032 binding to the kinase domains of wild-type BRAF and CRAF is required for increased RAF dimerization remains unclear. To address this, we generated two mutant CAF lines expressing gatekeeper mutations in the kinase domain of BRAF and CRAF that prevent access of BRAFis to their ATP-binding pockets: one wild-type BRAF-deficient line expressing BRAF(T529N) and another wild-type CRAF-deficient line expressing CRAF(T421N) (Fig. S5A, D)^40^. The depletion of endogenous BRAF or CRAF significantly reduced PLX4032-induced nuclear β-catenin accumulation in control CAFs (siBRAF and siCRAF). A similar nuclear β-catenin reduction was observed in CAFs expressing the gatekeeper mutants BRAF (siBRAF+T529N) and CRAF (siCRAF+T421N), but not in CAFs expressing wild-type BRAF (siBRAF+BRAF-WT) or CRAF (siCRAF+CRAF-WT) (Figs. S5B-C, E-F). Notably, PLX4032 and GSK2118436 failed to induce BRAF-CRAF heterodimer formation in CAFs expressing either BRAF (T529N) or CRAF (T421N) (Fig. S6). As expected, PLX4032 did not promote the phosphorylation of BRAF Ser445 or CRAF Ser338 in these gatekeeper mutant lines because of the inhibition of BRAF and CRAF dimerization (Fig. 5L, M). Together, these findings demonstrate that the direct binding of BRAFis to the kinase domains of BRAF or CRAF is a key step for RAF dimerization, activation, and downstream nuclear accumulation of β-catenin in CAFs.

### BRAFi-induced RAF dimerization and phosphorylation are RAS dependent

RAFs are mediators between RAS-GTPases and downstream MEK and ERK kinases. In wild-type RAF cells, RAS binding to the RAS-binding domain (RBD) and cysteine-rich domain (CRD) of RAF is essential for RAF activation because it effectively eliminates autoinhibition, releases the kinase domains, and facilitates RAF dimerization and transactivation. To determine whether RAS binding is required for BRAFi-induced RAF dimerization and transactivation, we depleted all RAS isoforms in CAFs using siRNA (Fig. 6A). PLA revealed that the PLX4032-induced increase in BRAF-CRAF heterodimer formation was markedly reduced in RAS-depleted CAFs compared with mock-transfected cells (Figs. 6B and S7A). Consistent with this, the phosphorylation of BRAF at Ser445 and CRAF at Ser338 was not elevated following PLX4032 treatment in RAS-depleted cells (Fig. 6C), confirming that RAS is involved in BRAFi-driven RAF dimerization and phosphorylation.

**Figure 6.**
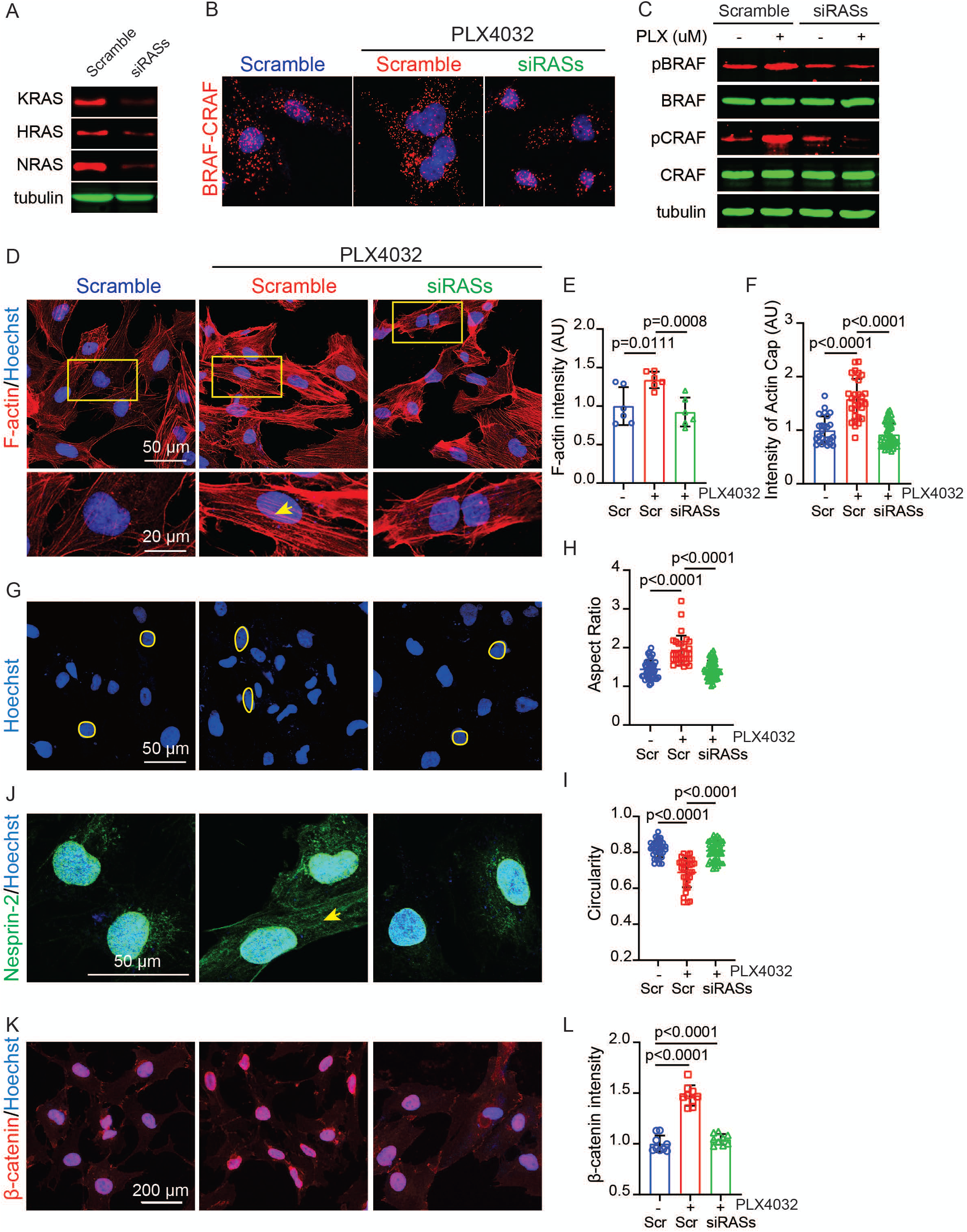
BRAFi-induced RAF activation is RAS dependent. (A) Western blot confirming effective silencing of KRAS, HRAS, and NRAS expression in iM27 using siRNAs. (B) PLA results showing BRAF and CRAF heterodimerization in iM27 transfected with scramble siRNA (Scr), scramble iM27 treaded with PLX4032, and RAS-deficient iM27 (siRASs) treated with PLX4032. (C) Western blot showing the phosphorylation of BRAF (Ser445) and CRAF (Ser338) in iM27 transfected with scramble siRNA and RAS-deficient iM27 with and without PLX4032 treatment. (D) Confocal images of F-actin expression and organization in iM27 cells transfected with scramble siRNA, scramble iM27 cells treated with PLX4032 (scramble), and RAS-deficient iM27 cells (siRASs) treated with PLX4032. The insets display enlarged views of representative individual cells (highlighted by yellow boxes). Yellow arrows indicate actin caps. (E–F) Quantification of F-actin intensity (F) and actin cap intensity (F) using ImageJ. The data are presented as the mean ± SD (n = 6 randomly selected 20 x fields per group for E; n = 25-52 nuclei per group for F). (G) Confocal images of nuclei visualized by Hoechst staining in iM27 transfected with scramble siRNA, scramble iM27 treaded with PLX4032, and RAS-deficient iM27 (siRASs) treated with PLX4032. The nuclear boundaries are outlined with yellow circles. (H–I) Quantification of nuclear morphology based on the confocal images shown in (G), including the nuclear aspect ratio (H) and circularity (I), was performed using ImageJ. The data are presented as the mean ± SD (n = 30-67 nuclei per group). (J) Confocal images showing Nesprin-2 distribution in iM27 cells transfected with scramble siRNA, scramble iM27 cells treated with PLX4032, and RAS-deficient iM27 cells (siRASs) treated with PLX4032. The yellow arrow indicates the abnormal cytosolic localization of Nesprin-2. (K) Representative fluorescence images showing nuclear β-catenin staining in iM27 cells transfected with scramble siRNA, scramble iM27 treaded with PLX4032, and RAS-deficient iM27 (siRASs) treated with PLX4032. (L) Quantification of nuclear β-catenin intensity under the indicated conditions shown in (K). The data are presented as mean ± SD (n = 9 randomly selected 20X fields per group).

We next examined whether disruption of the RAS-RAF signaling axis affects cytoskeletal remodeling, nuclear morphology, and β-catenin nuclear accumulation. Confocal imaging of F-actin expression and distribution revealed that PLX4032-induced actin polymerization and actin cap formation were suppressed by RAS depletion (Figs. 6D-F and S8A-C). Similarly, RAS-depleted CAFs reversed PLX4032-induced nuclear deformation, with the nuclear aspect ratio returning to ∼1.44 and the circularity increasing to ∼0.81 (Figs. 6G-I and S8D-F). In addition, the abnormal cytosolic redistribution of Nesprin-2 observed after PLX4032 treatment was reversed in RAS-depleted cells (Figs. 6J and S8G). As a result, the increased nuclear accumulation of β-catenin induced by PLX4032 was significantly diminished in RAS-depleted CAFs, as measured by the average intensity of nuclear β-catenin (Figs. 6K-L and S8H-I). These findings confirmed that RAS binding to wild-type BRAF and CRAF and releasing the kinase domains are essential for BRAFi-induced nuclear deformation and β-catenin accumulation in CAFs.

### ROCK signaling regulates cytoskeletal reorganization and β-catenin nuclear translocation

To determine how RAF activation promotes actin fiber polymerization, we focused on the ROCK signaling pathway, a key regulator of the actin cytoskeleton. Notably, we observed the activation of the ROCK pathway following PLX4032 treatment. MYPT1, a well-known substrate of ROCK that regulates actin contraction and relaxation, was phosphorylated upon PLX4032 exposure (Fig. 7A). Phosphorylation of MYPT1 inhibits its activity, thereby maintaining MLC phosphorylation and promoting actin filament polymerization and contraction^41^. These results suggest that β-catenin translocation to the nucleus is a ROCK-dependent process.

**Figure 7.**
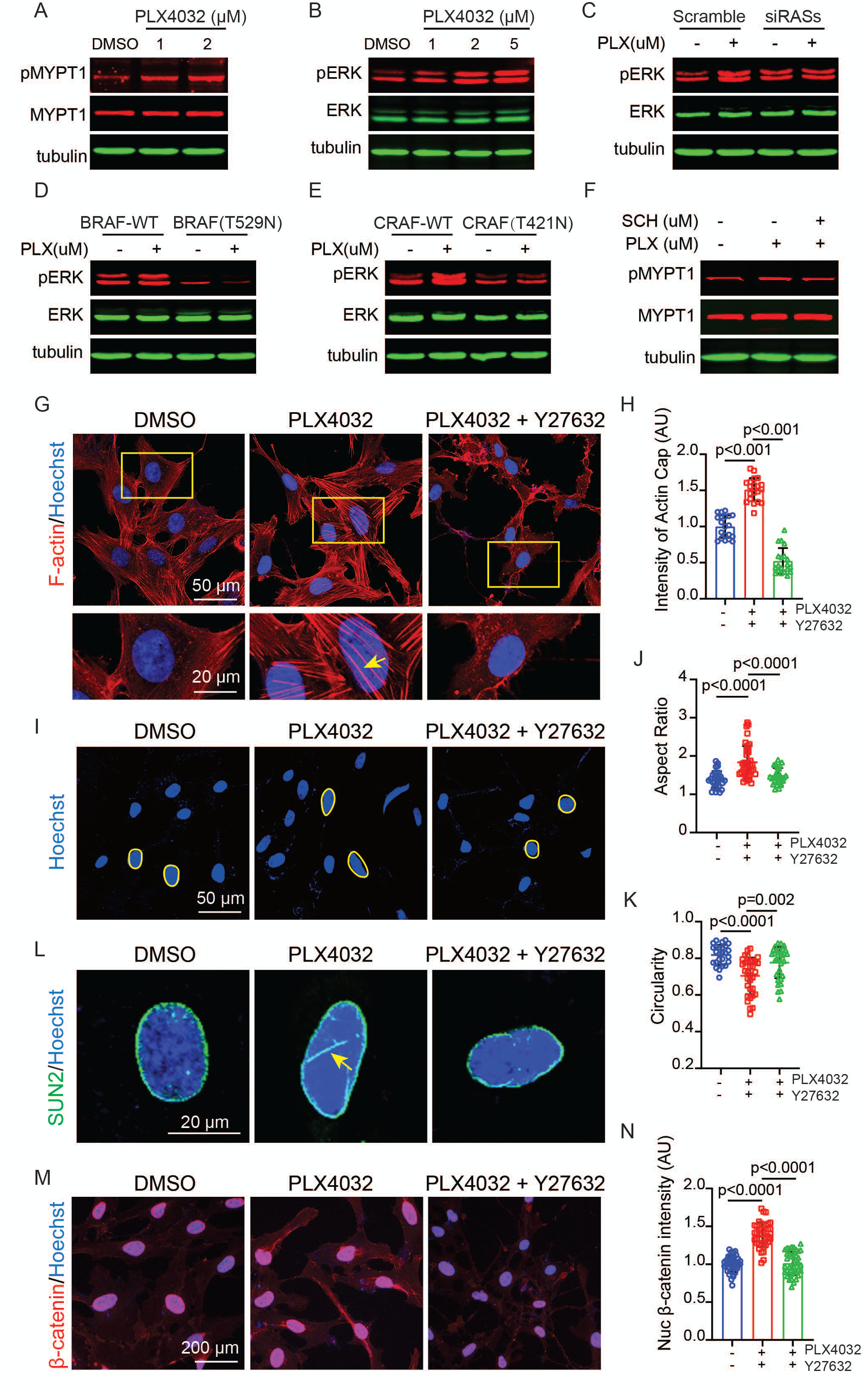
ROCK signaling is downstream of RAF activation to promote cytoskeletal remodeling and nuclear deformation. (A) Western blot showing increased phosphorylation of MYPT1 in PLX4032-treated iM27. (B) Western blot showing increased ERK phosphorylation in response to increasing concentrations of PLX4032 in iM27 cells. (C) Western blot showing the phosphorylation of ERK in iM27 transfected with scramble siRNA and RAS-deficient iM27 (siRASs) with and without PLX4032 treatment. (D–E) Western blots showing ERK phosphorylation in BRAF-deficient iM27 cells overexpressing wild-type BRAF and BRAF(T529N) (D) and in CRAF-deficient iM27 cells overexpressing wild-type CRAF and CRAF(T421N) (E) with and without PLX4032 treatment. (F) Western blot showing the phosphorylation of MYPT1 in iM27 cells treated with DMSO, PLX4032, or a combination of PLX4032 and the ERK inhibitor SCH772984 (SCH). (G) Confocal images of F-actin expression and organization in iM27 cells treated with DMSO, PLX4032, or a combination of PLX4032 and the ROCK inhibitor Y27632. The insets display enlarged views of representative individual cells (highlighted by yellow boxes). Yellow arrows indicate the actin caps. (H) Quantification of the actin cap intensity via ImageJ shown in (G). Data are presented as mean ± SD (n = 20 nuclei per group). (I) Representative confocal images of nuclear morphology visualized by Hoechst staining in iM27 cells treated with DMSO, PLX4032, and a combination of PLX4032 and the ROCK inhibitor Y27632. (J–K) Quantification of nuclear morphology based on the confocal images shown in (I), including the nuclear aspect ratio (J) and circularity (K), was performed via ImageJ. The data are presented as mean ± SD (n = 27-38 nuclei per group). (L) Confocal images showing SUN2 distribution in iM27 cells treated with DMSO, PLX4032, and a combination of PLX4032 and the ROCK inhibitor Y27632. The yellow arrow indicates the abnormal nuclear localization of SUN2 in PLX4032-treated cells. (M) Representative fluorescence images showing nuclear β-catenin staining in iM27 cells treated with DMSO, PLX4032, or a combination of PLX4032 and the ROCK inhibitor Y27632. (N) Quantification of the nuclear β-catenin intensity under the indicated conditions shown in (M). Data are presented as mean ± SD (n = 40 nuclei per group).

To assess possible molecular interactions between the RAS-RAF signaling pathway and ROCK signaling, we first evaluated the status of ERK, which is an established downstream effector of RAF kinases^42^, in response to increasing PLX4032 and GSK2118436 concentrations in the two CAF lines. The results showed that BRAFi treatment increased ERK phosphorylation (Figs. 7B and S9A-C). This response was diminished upon RAS depletion (Figs. 7C and S9D), confirming that RAS is the upstream activator of the BRAFi-induced RAF-ERK pathway. Furthermore, PLX4032 did not promote ERK phosphorylation in cells expressing BRAFi-resistant BRAF (T529N) or CRAF (T421N) mutants, indicating that BRAFi-driven RAF activation is necessary for activation of downstream ERK signaling (Fig. 7D, E). Downstream MYPT1 Phosphorylation induced by PLX4032 was diminished by cotreatment with the ERK inhibitor SCH772984 (SCH) (Fig. 7F), further confirming that ROCK acts downstream of the RAF-ERK pathway to promote actin polymerization. In addition, GSK-3β phosphorylation at Serine 9 was significantly increased in CAFs following PLX4032 treatment (Fig. S9E), indicating that GSK-3β inactivation contributed to nuclear β-catenin accumulation. Immunostaining for β-catenin also showed that the PLX4032-induced increase in nuclear β-catenin was reduced by ERK inhibition (Fig. S9F, G).

Next, we investigated whether the inhibition of ROCK signaling could reverse the effects of PLX4032 on nuclear deformation and β-catenin nuclear accumulation in CAFs. To this end, CAFs were cotreated with PLX4032 and a well-established ROCK inhibitor Y27632. Compared with PLX4032 alone, F-actin staining confirmed effective actin cytoskeletal disruption and diminished actin cap formation under combined treatment (Fig. 7G, H). This observation confirms that PLX4032-induced actin cap formation depends on the activation of the ROCK pathway.

Furthermore, PLX4032-induced nuclear deformation was reversed by Y27632 treatment, as demonstrated by the restoration of nuclear morphology, with the nuclear aspect ratio returning to ∼1.50 and the circularity increasing to ∼0.78 (Fig. 7I-K). The aberrant nuclear distribution of SUN2 triggered by PLX4032 was rescued in CAFs treated with Y27632 (Fig. 7L). As expected, increased nuclear accumulation of β-catenin induced by PLX4032 was significantly reduced by ROCK inhibition (Fig. 7M, N). The effects of Y27632 in reversing PLX4032-induced nuclear deformation, actin polymerization, and nuclear β-catenin accumulation were confirmed using another ROCK inhibitor, HA-1077, and in additional CAF line (Fig. S10).

Because increased ECM stiffness triggered actin polymerization, nuclear deformation, and nuclear β-catenin import in CAFs, we wanted to confirm that inhibiting ROCK activity has the similar suppressive effects on CAFs cultured on a hard substrate as PLX4032-treated CAFs. CAFs seeded on hard substrate with a stiffness of 50 kPa were treated with Y27632 or HA-1077. upon ROCK inhibition, confocal imaging revealed a remarkable loss of cytoskeletal actin fibers and actin cap structures (Figs. 8A, B and S11A-D). Consequently, nuclear deformation was reversed with the nuclear aspect ratio reduced to ∼1.55 and the circularity increased to ∼0.78 (Figs. 8C-E and S11E-G). The aberrant nuclear distribution of SUN2 induced by stiffness was reduced in the Y27632-treated CAFs (Fig. 8F). Importantly, the increased nuclear translocation of β-catenin observed in CAFs cultured on stiff substrates was markedly diminished following Y27632 or HA-1077 treatment (Figs. 8G, H and S11H-K). Collectively, these findings indicated that the ROCK pathway is a central mediator that enables CAFs to respond to BRAFi and stiffness, reinforcing their activation through cytoskeletal remodeling, nuclear deformation, and β-catenin nuclear accumulation.

**Figure 8.**
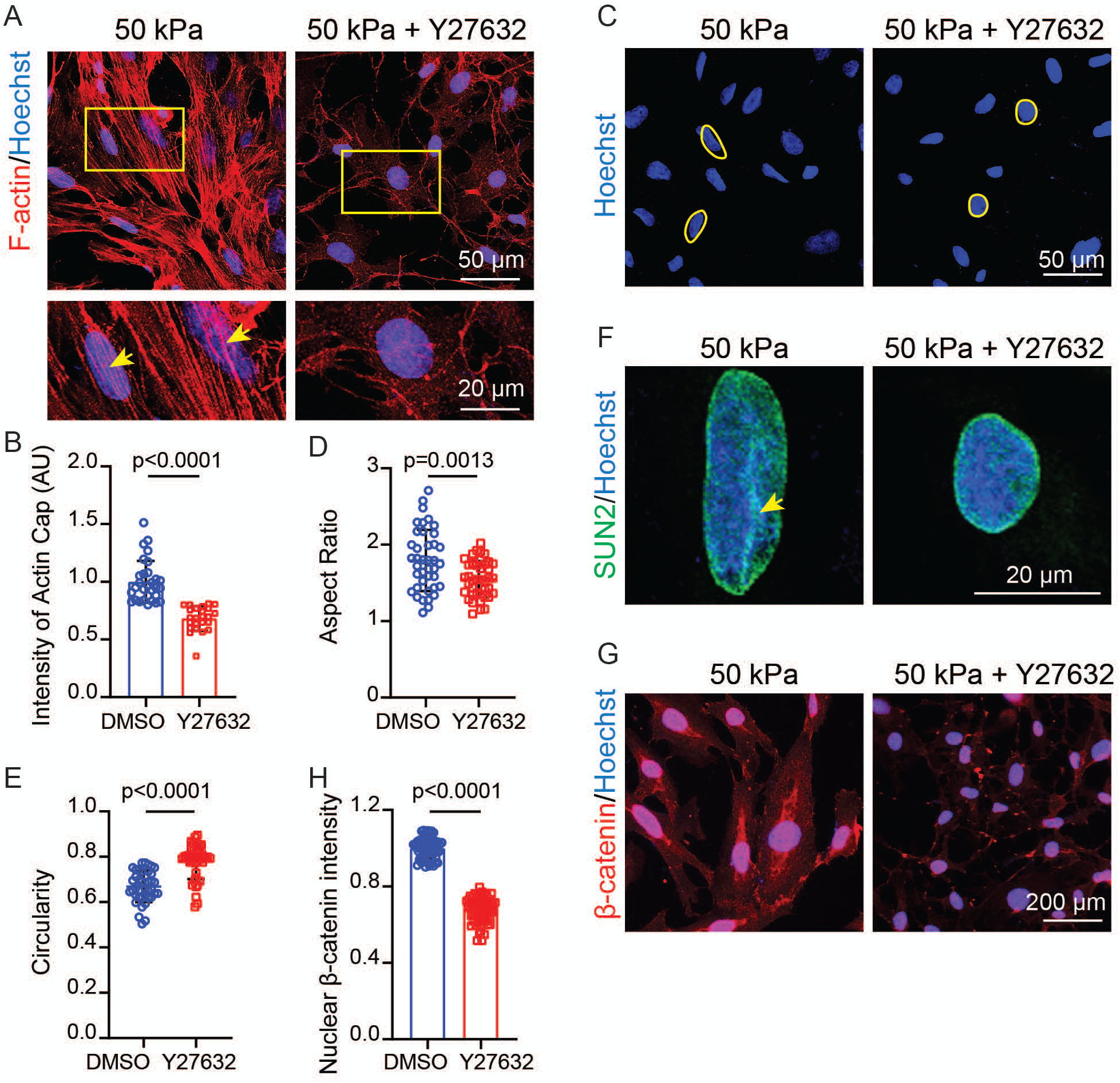
ROCK inhibitor reverses stiffness-induced nuclear deformation and β-catenin accumulation. (A) Confocal images of F-actin expression and organization in iM27 cells cultured on hard slides with a stiffness of 50 kPa with or without ROCK inhibitor Y27632 treatment. The representative individual cells shown in the insets correspond to the cells marked with yellow boxes in the images above. The actin caps are indicated by yellow arrows. (B) Quantification of the actin cap intensity using ImageJ, as shown in (A). The data are presented as mean ± SD (n = 24-30 nuclei per group). (C) Representative confocal images showing nuclei stained with Hoechst in iM27 cells cultured on hard slides with a stiffness of 50 kPa with or without ROCK inhibitor Y27632 treatment. Yellow circles highlight the approximate nuclear boundaries. (D–E) Scatter dot plots showing nuclear morphological changes under the indicated conditions corresponding to (C). The nuclear aspect ratio (D) and circularity (E) were analyzed and quantified using ImageJ. Data are presented as mean ± SD (n = 40 nuclei per group). (F) Confocal images showing SUN2 staining in iM27 cells cultured on hard slides with a stiffness of 50 kPa with or without ROCK inhibitor Y27632 treatment. Disorganized SUN2 distribution in iM27 cells cultured on 50 kPa slides is indicated by yellow arrows. (G) Fluorescence microscopy images showing nuclear β-catenin expression in iM27 cells cultured on hard slides with a stiffness of 50 kPa with or without ROCK inhibitor Y27632 treatment. (H) Quantification of nuclear β-catenin intensity in iM27 cells corresponding to (G). Data are presented as mean ± SD (n = 60 cells per group).

## DISCUSSION

In this study, we identified a shared cytoskeleton-to-nucleus signaling mechanism by which CAFs respond to BRAF inhibition and matrix stiffness and reinforce their activation. Mechanistically, BRAFis activate the RAF-ROCK axis, inducing actin cytoskeletal remodeling and nuclear deformation in CAFs. This process disrupts nuclear membrane integrity and facilitates β-catenin accumulation in the nucleus, a central mediator of Wnt/β-catenin signaling, which not only regulates transcriptional programs linked to cell fate, tissue remodeling, and cancer progression, and further promotes actin fiber polymerization, establishing a cytoskeletal-nuclear feedback loop. We demonstrated that β-catenin is a functional driver of CAF activation in melanoma both *in vitro* and *in vivo*. This reinforcing cycle between actin polymerization and nuclear β-catenin defines a critical regulatory axis that amplifies signaling and sustains cellular plasticity, enabling CAF adaptation to mechanical and biochemical cues.

The nucleus, enclosed by a double-layered membrane, tightly regulates protein transport through nuclear pore complexes (NPCs)^43^. Despite lacking a canonical nuclear localization signal, β-catenin may enter the nucleus via two routes: direct NPC interaction or piggyback transport with nuclear proteins, such as TCF/LEF^44,45^. While both mechanisms may operate in BRAFi-treated CAFs, our data support an alternative mechanism through which BRAFi-induced nuclear deformation alters NPC architecture and enhances β-catenin nuclear import. Additionally, we found that BRAFis promoted GSK-3β phosphorylation, further increasing β-catenin stabilization and contributing to its nuclear presence^46^. Increased stiffness is a common characteristic of metastatic tumors. Nuclear shape and NPC function are known to be regulated by ECM stiffness and mechanical forces^47,48^. We showed that exposure to a stiff matrix induces nuclear deformation and nuclear β-catenin accumulation in CAFs as well. Collectively, these findings indicate that mechanically tuned nuclear signaling enables CAFs to sense and adapt to external pressures, thereby promoting their pro-tumorigenic functions.

Nuclear morphology exhibits substantial variability and is shaped by mechanical forces, cytoskeletal tension, molecular signaling, and cellular context to maintain nuclear function^49^. Nuclear pores are large multiprotein structures within the nuclear membrane that transport molecules between the cytoplasm and nucleus^50^. Thus, changes in the nuclear shape and nuclear membrane tension can alter the diameter of the nuclear pores. In both BRAFi-treated and stiffness-stimulated CAFs, we observed nuclear elongation and NPC disruption, conditions that increased nuclear import of β-catenin (Fig. 9). This mechanically induced nuclear access allows CAFs to integrate diverse environmental stimuli into unified cellular and transcriptional responses.

**Figure 9.**
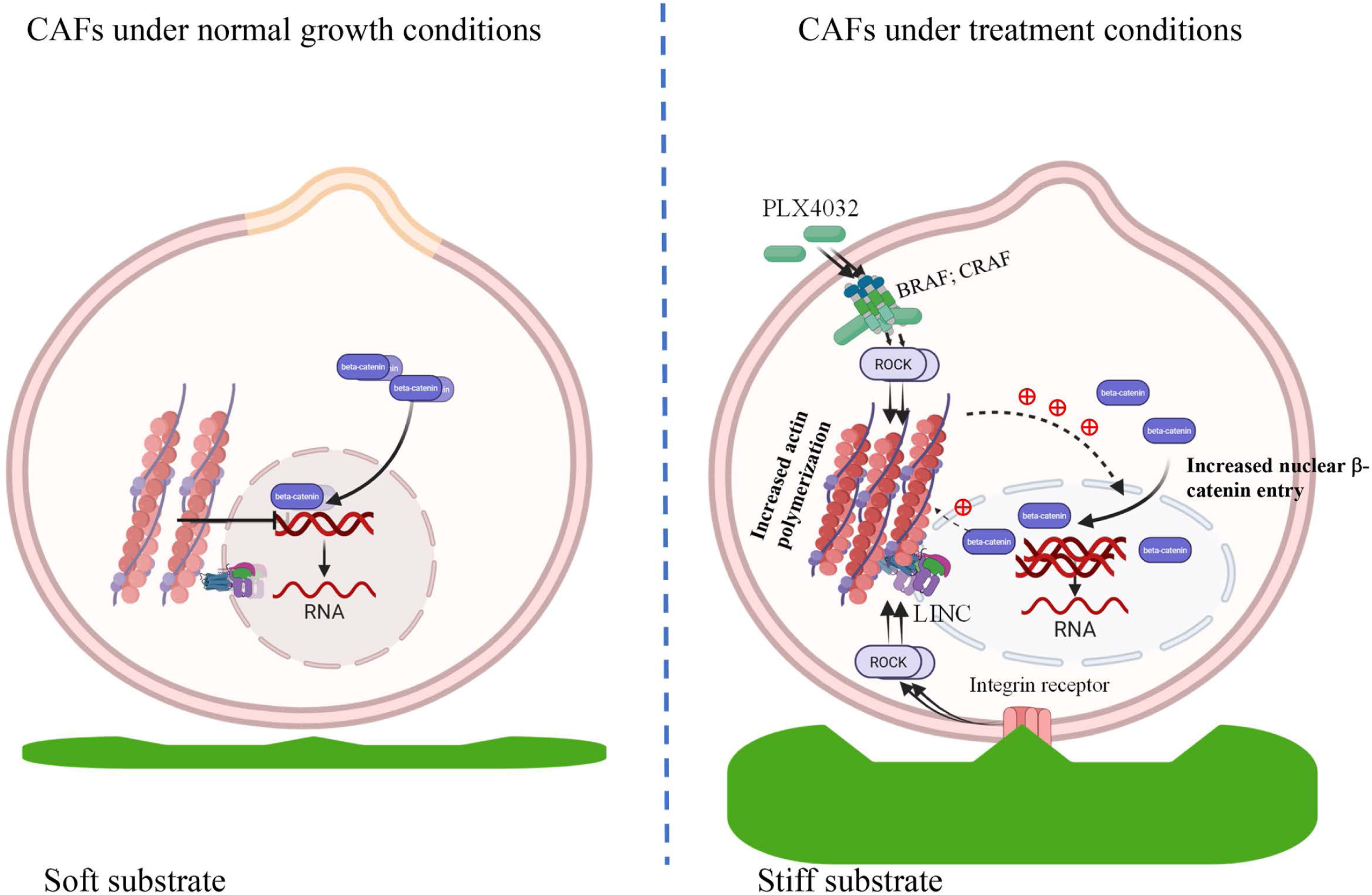
A unique response mechanism by which CAFs respond to various external stimuli and reinforce their activation via a positive feedback loop. The ROCK-cytoskeleton axis can be activated in CAFs by BRAFis and stiff substrates and drives nuclear deformation, which subsequently promotes increased β-catenin nuclear translocation, which further drives actin stress fiber polymerization, forming a reciprocal, feed-forward loop to continuously reinforce CAF functions. Created in BioRender. Zhang, Y. (2026) https://BioRender.com/kyqkmtp

The mechanisms through which both BRAFis and a stiff environment lead to nuclear deformation are intriguing. Transmembrane LINC complexes mechanically connect lamins and chromatin inside the nucleus to the cytoskeleton outside the nucleus^51^. This connection is essential for maintaining the nuclear shape and structure and for transmitting mechanical forces inside the nucleus. Furthermore, actin polymerization drives the formation of actin caps, which specifically influences nuclear shape and positioning^35^. We consistently observed increased actin polymerization and actin cap presence, disorganized SUN2 and Nesprin-2 localization, and elongated nuclei in BRAFi-treated CAFs and CAFs cultured on hard substrates, suggesting that actin polymerization driven by BRAFis and increased matrix stiffness exerted mechanical forces to deform the nucleus through disruption of the LINC complex. However, the exact mechanism underlying the changes in nuclear membrane integrity that increase β-catenin nuclear entry remains to be further elucidated.

The rearrangement and assembly of the actin cytoskeleton not only control cell shape, polarity, and migration but also influence cell metabolism^52^. Increased actin polymerization was observed in CAFs exposed to either BRAFis or stiff matrix, although the stimulatory mechanisms may differ (Fig. 9). It is known that actin polymerization is regulated by the ROCK signaling pathway^53^. While matrix stiffness can activate ROCK signaling through integrin receptors on the cell membrane^54^, our findings reveal a novel signaling mechanism through which BRAFis induces abnormal activation of RAF kinases in CAFs, leading to the activation of the ROCK signaling pathway. We confirmed that ERK signaling functions downstream of the RAS/RAF axis in CAFs upon BRAFi treatment. The activation of ERK signaling has multiple effects, as it can activate the ROCK pathway and inactivate GSK-3β via phosphorylation. These data further demonstrate that CAFs integrate diverse extracellular cues to reprogram their functional state through ROCK-dependent cytoskeletal remodeling and nuclear deformation.

Although BRAFi was designed to inhibit mutant BRAF carrying the V600E point mutation by inserting in the mutant kinase domain and blocking the active site, it has been shown that it can paradoxically activate the MAPK/ERK pathway in cells carrying wild-type BRAF. However, how CAFs react to BRAFis and change their behavior in melanoma remains unknown. Our data revealed that both BRAF and CRAF function in BRAF-wild-type CAFs by accelerating BRAF and CRAF dimerization and phosphorylation in CAFs through RAS-dependent and kinase domain-mediated mechanisms. Another potential promoting mechanism is that BRAFis in wild-type BRAF CAFs can stabilize BRAF and CRAF dimers. Theoretically, increased RAF activity explains the observation that the ERK and ROCK pathways are activated in CAFs by BRAFis, contributing to increased actin polymerization, which acts on the nucleus.

Our data showed that BRAFis can bind in the wild-type kinase domain although it was designed to fit in the mutant BRAF kinase domain and block its abnormal catalytic activity. The opposite effects of BRAFis, promoting wild-type BRAF kinase activity while inhibiting the mutant BRAF kinase, can be attributed to the BRAF mutation and the steric conformation of the kinase domains. As shown in Fig. 10, in wild-type monomeric BRAF, binding of the N-terminal RBD and CRD to the C-terminal kinase domain blocks the access of kinase to either kinase substrates or small-molecule compounds, such as BRAFis, thereby BRAF remains inactive. However, in the BRAF V600E mutant, this specific substitution within the activation loop in the kinase domain changes the steric conformation and disrupts autoinhibition, leading to its constitutive activation even as a monomer without RAS binding. BRAFis can effectively access the V600E mutant kinase domain and inhibit its kinase activity. In BRAF-wild-type CAFs, BRAFi can only enter the kinase domains, which do not contain the V600E mutation, in BRAF and CRAF after RAS binds to the RBD and CRD domains and release the autoinhibition. Surprisingly, BRAFis then induces unexpected conformational changes in the wild-type kinase domains and promotes RAF dimerization and transactivation instead of inhibition.

**Figure 10.**
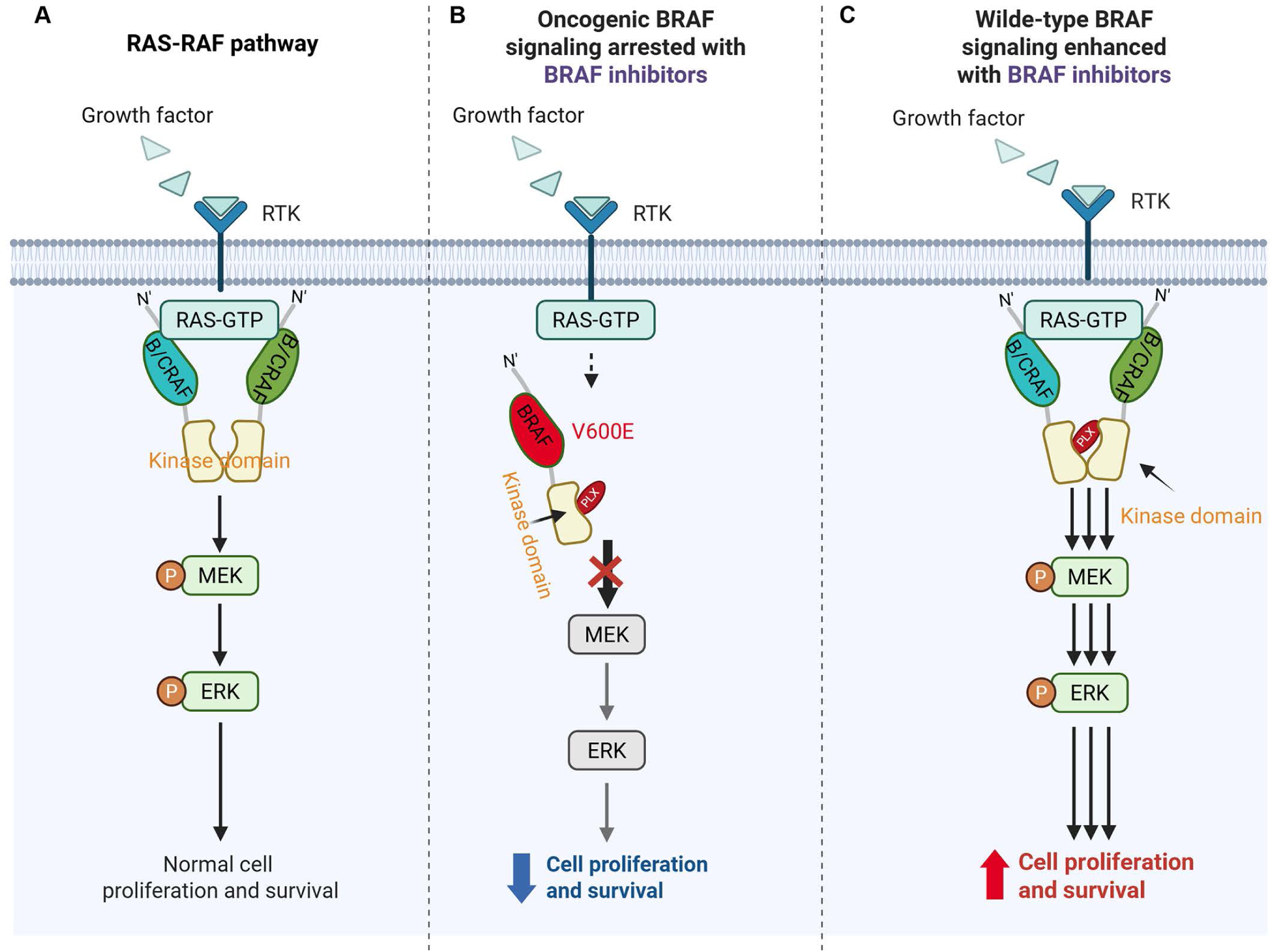
BRAFi accelerates BRAF and CRAF dimerization and transactivation in CAFs. (A) In BRAF wild-type cells, without BRAFi treatment, RAF kinases are activated when a cell receives signals from its environment, usually through receptor tyrosine kinases (RTKs), and activate the small GTPase RAS. Activated RAS binds to BRAF and CRAF and releases autoinhibition, leading to RAF transactivation and the activation of downstream MEK and ERK. (B) In BRAF-mutant cells, BRAF possesses a mutant but active kinase domain that can activate downstream MEK and ERK without the need of RAS binding. BRAFis are designed to enter the mutant kinase domain and blocks its activity. (C) In BRAF wild-type CAFs, BRAF and CRAF need RAS binding to release the kinase domain, which do not carry V600E mutation. Once BRAFis gain the access to the kinase domains of BRAF and CRAF, they can induce unexpected conformational changes, which accelerate BRAF and CRAF dimerization and transactivation, leading to increased downstream MEK and ERK signaling activities. Created in BioRender. Siegfried, L. (2026) https://BioRender.com/ie7p9ak.

Given the role of CAFs in promoting melanoma progression and therapy resistance, they represent attractive but elusive therapeutic targets. Our findings uncover a common mechanotransduction pathway by which CAFs sense and integrate chemical and mechanical cues via ROCK– cytoskeletal signaling. This axis regulates nuclear shape and allows mechanoresponsive transcription factors such as β-catenin to enter the nucleus and rewire CAF function. Inhibition of the ROCK pathway suppressed CAF responses to both BRAFis and ECM stiffness and blocked transcription factor nuclear entry. Taken together, these results identify the ROCK–actin–nuclear axis as a central regulator of CAF plasticity and a promising target for stroma-directed therapies for melanoma or other types of solid tumors.

## ETHICAL APPROVAL

The Institutional Animal Care and Use Committee of the University of Cincinnati (IACUC) approved all experimental procedures involving mice under protocol 22-08-19-01.

## ACKNOWLEDGMENTS

We thank Dr. Dorothy Supp for providing human dermal fibroblasts. This work was supported by the NIH R01 CA249737 (YZ), and the Center for Clinical & Translational Science & Training (CCTST) Just-In-Time Core Grant (YZ), the UCCC Pilot Project Award Program (YZ), and the Marlene Harris Ride Cincinnati Cancer Pilot Program (YZ). The CCTST Just-In-Time Core Grant was supported by the National Center for Advancing Translational Sciences of the National Institutes of Health under Award Number UL1TR001425. The content is the sole responsibility of the authors and does not necessarily represent the official views of the NIH.

## AUTHOR CONTRIBUTIONS

Y.Z. designed the experiments; J.W., B.S., Y.X. and L.Z. performed the experiments and collected the data; J.W., B.S., Y.X., L.Z. and Y.Z. analyzed the data; S.E.M and M.X. provided the mouse models; J.W., Y.Z. and T.A. wrote the main manuscript text; J.W. and Y.Z. prepared the figures; J.W., L.Z., Y.Z., L.S., S.E.M. and T.A. reviewed and edited the manuscript. All authors have reviewed and approved the final version of the manuscript. All authors agree to be accountable for all aspects of the work in ensuring that questions related to the accuracy or integrity of any part of the work are appropriately investigated and resolved and declare confidence in the integrity of the contributions of their coauthors.

## CONFLICTS OF INTEREST

The authors declare that they have no competing interests.

## Notes

### Competing Interest Statement

The authors have declared no competing interest.

### Summary of Updates

The manuscript was reorganized to provide a better flow, otherwise no any significant changes were made from previous version.

