## Supplementary figures for "Nuclear force transmission drives cancer-associated fibroblast activation under BRAF inhibition"

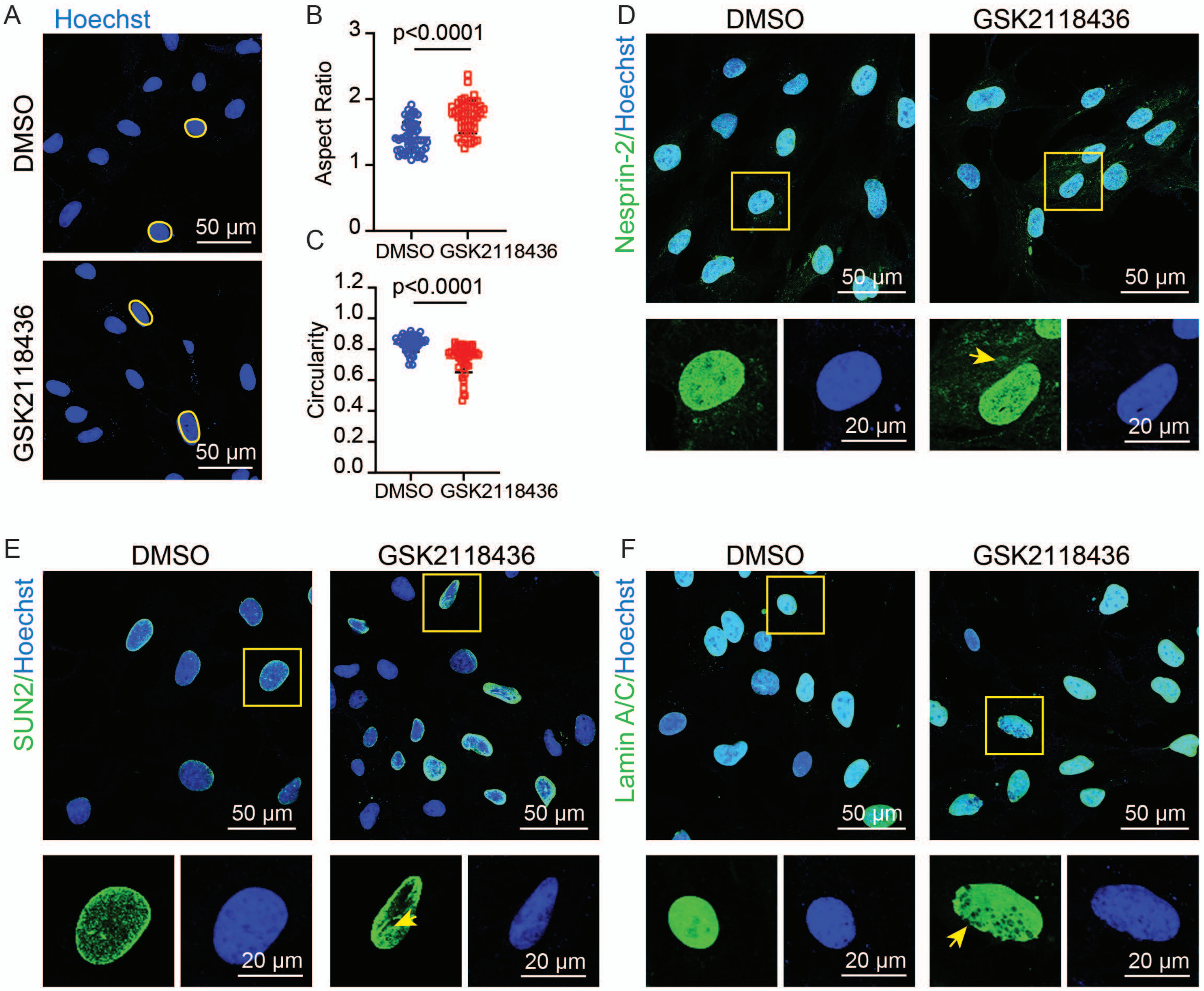

A
Hoechst
DMSO
GSK2118436
50 μm
50 μm
B
p<0.0001
Aspect Ratio
3
2
1
0
DMSO
GSK2118436
C
p<0.0001
Circularity
1.2
1.0
0.8
0.6
0.4
0.2
0.0
DMSO
GSK2118436
D
DMSO
GSK2118436
Nesprin-2/Hoechst
50 μm
50 μm
20 μm
20 μm
E
DMSO
GSK2118436
SUN2/Hoechst
50 μm
50 μm
20 μm
20 μm
F
DMSO
GSK2118436
Lamin A/C/Hoechst
50 μm
50 μm
20 μm
20 μm


**Fig. S1**

**Figure S1 BRAFi induce nuclear deformation in CAFs.**

(A) Representative confocal images showing nuclei stained with Hoechst in iM27 w/ or w/o GSK2118436 treatment. Yellow circles highlight the approximate nuclear boundaries. iM27 cells were treated with either DMSO or GSK2118436 for comparison.

(B–C) Scatter dot plots showing nuclear morphological parameters in iM27 w/ or w/o GSK2118436 treatment. (B) the nuclear aspect ratio and (C) circularity were analyzed and quantified from confocal images using ImageJ. The data are presented as mean ± SD (n = 47-56 nuclei per group).

(D–F) Confocal images showing immunostaining of key components of the LINC complex in iM27 w/ or w/o GSK2118436 treatment, including (D) Nesprin-2, (E) SUN2, and (F) nuclear Lamin A/C. In each figure, individual cells shown in small boxes are representative stained cells denoted by yellow boxes in the large boxes above. Disrupted distribution patterns of Nesprin-2, SUN2, and Lamin A/C in GSK2118436-treated iM27 cells are indicated by yellow arrows.

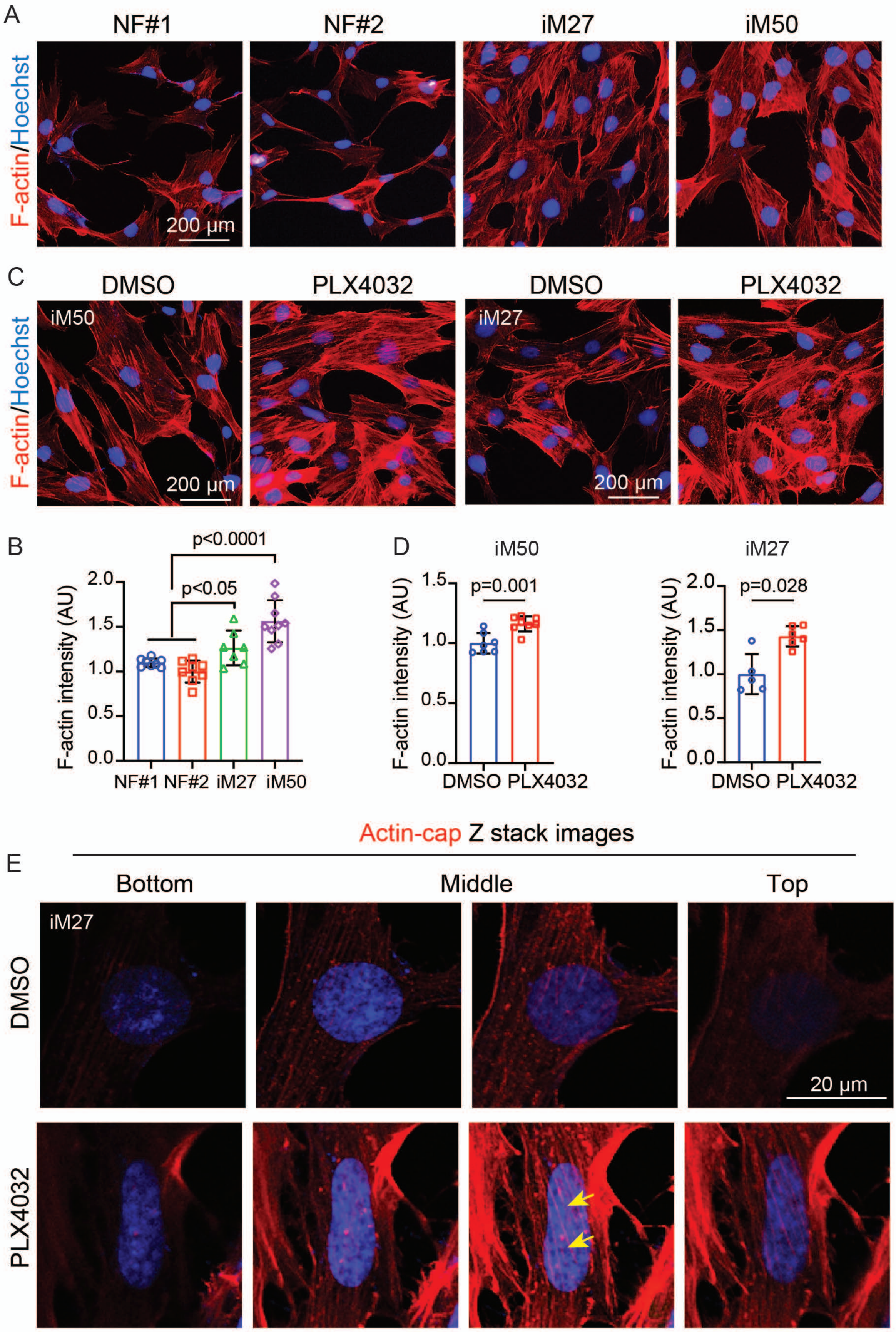

A
NF#1
NF#2
iM27
iM50
F-actin/Hoechst
200 µm
C
DMSO
PLX4032
DMSO
PLX4032
iM50
iM27
F-actin/Hoechst
200 µm
200 µm
B
p<0.0001
p<0.05
F-actin intensity (AU)
2.0
1.5
1.0
0.5
0.0
NF#1 NF#2 iM27 iM50
D
iM50
p=0.001
F-actin intensity (AU)
1.5
1.0
0.5
0.0
DMSO PLX4032
iM27
p=0.028
F-actin intensity (AU)
2.0
1.5
1.0
0.5
0.0
DMSO PLX4032
Actin-cap Z stack images
E
Bottom
Middle
Top
iM27
DMSO
PLX4032
20 µm


Fig. S2

**Figure S2 BRAFi induces actin polymerization and actin cap formation in CAFs.**

(A) Representative fluorescence images showing F-actin expression and organization in normal human fibroblasts (NF#1 and NF#2) and CAFs (iM27 and iM50).

(B) Quantification of F-actin intensity in normal human fibroblasts and CAFs. The data are presented as mean ± SD (n = 7-9 random 20X fields).

(C) Representative fluorescence images showing F-actin expression and organization in DMSO-treated and PLX4032-treated iM50 and iM27 CAFs.

(D) Quantification of F-actin intensity in DMSO-treated and PLX4032-treated iM50 and iM27 CAFs. The data are presented as mean ± SD (n = 7-8 random 20X fields).

(E) Representative Z-stack confocal images of F-actin in iM27 cells treated with DMSO or PLX4032. A single nucleus was scanned from bottom to top at 1 μm intervals. Actin caps spanning across the Hoechst-stained nucleus are indicated by yellow arrows.

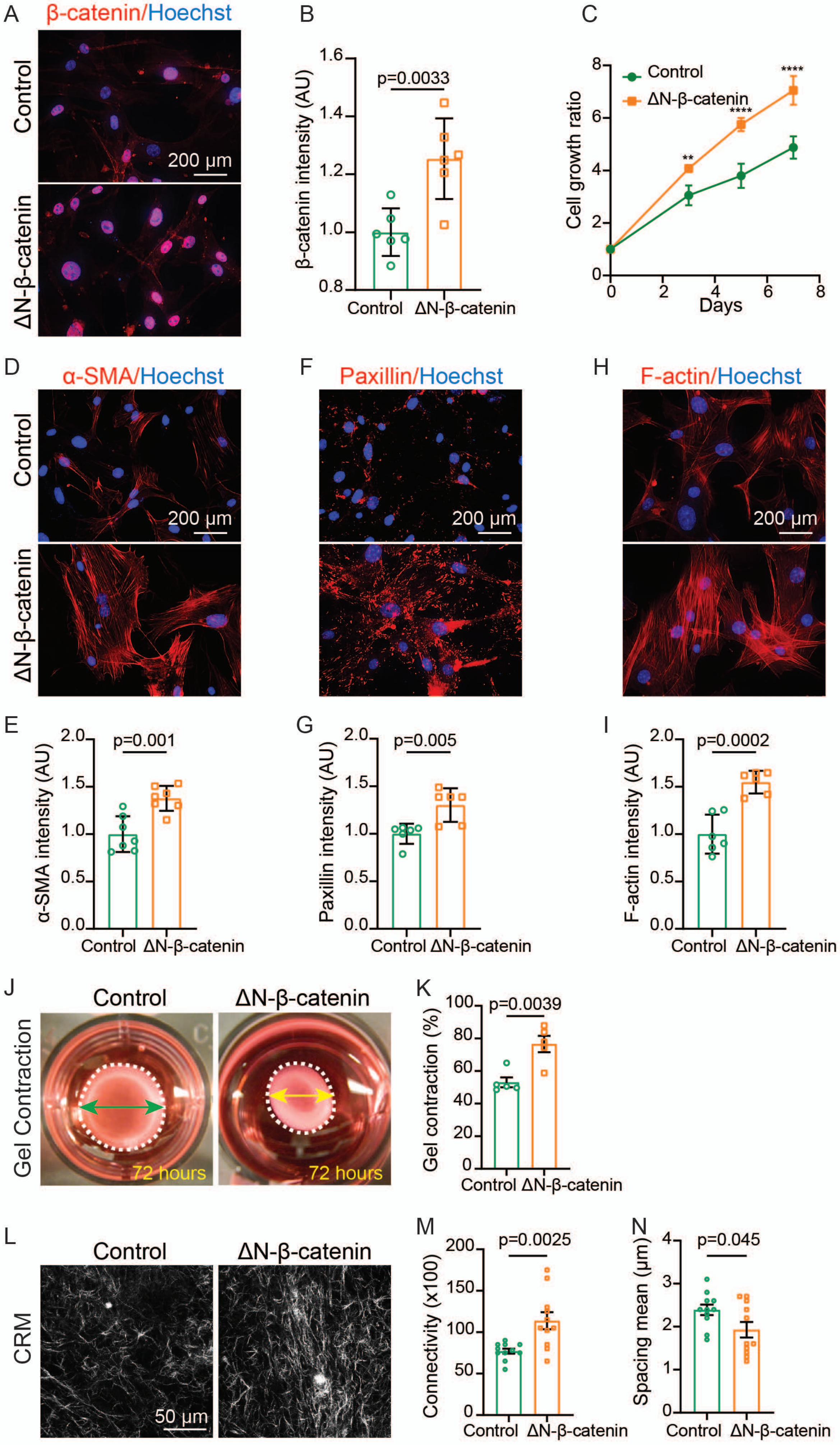
A
β-catenin/Hoechst
Control
ΔN-β-catenin
200 μm
B
p=0.0033
β-catenin intensity (AU)
1.6
1.4
1.2
1.0
0.8
Control
ΔN-β-catenin
C
Control
ΔN-β-catenin
Cell growth ratio
**
****
****
Days
D
α-SMA/Hoechst
F
Paxillin/Hoechst
H
F-actin/Hoechst
E
α-SMA intensity (AU)
p=0.001
G
Paxillin intensity (AU)
p=0.005
I
F-actin intensity (AU)
p=0.0002
J
Gel Contraction
72 hours
K
Gel contraction (%)
p=0.0039
L
CRM
50 μm
M
Connectivity (x100)
p=0.0025
N
Spacing mean (μm)
p=0.045

**Fig. S3**

**Figure S3 Nuclear β-catenin accumulation enhances the biological properties of fibroblasts.**

(A) Representative immunofluorescence images showing β-catenin expression in control fibroblasts (*Col1α2-CreER; Rosa-rtTA*) and β-catenin-overexpressing fibroblasts (*Col1α2-CreER; Rosa-rtTA; TetO-ΔN-β-catenin*). The nuclei were counterstained with Hoechst (blue).

(B) Quantification of nuclear β-catenin levels in control and ΔN-β-catenin-overexpressing fibroblasts. The data are presented as mean ± SD (n = 6 randomly selected 40X fields per group).

(C) Comparison of cell proliferation between control and ΔN-β-catenin-overexpressing fibroblasts. The data are presented as mean ± SD (n = 4 replicates per group).

(D) Representative immunofluorescence images showing α-SMA expression in control and ΔN-β-catenin-overexpressing fibroblasts. Nuclei were counterstained with Hoechst.

(E) Quantification of α-SMA expression in control and ΔN-β-catenin-overexpressing fibroblasts. The data are presented as mean ± SD (n = 6-7 randomly selected 40X fields per group).

(F) Representative immunofluorescence images showing paxillin expression in control and ΔN-β-catenin-overexpressing fibroblasts. Nuclei were counterstained with Hoechst.

(G) Quantification of paxillin expression levels in control and ΔN-β-catenin-overexpressing fibroblasts. The data are presented as mean ± SD (n = 6-7 randomly selected 40X fields per group).

(H) Representative immunofluorescence images showing F-actin expression in control and ΔN-β-catenin-overexpressing fibroblasts. Nuclei were counterstained with Hoechst.

(I) Quantification of F-actin levels in control and ΔN-β-catenin-overexpressing fibroblasts. The data are presented as mean ± SD (n = 6-7 randomly selected 40X fields per group).

(J) Representative images of collagen gel contraction assays for control and ΔN-β-catenin-overexpressing fibroblasts. The gel in each well is circled by a white dashed line. The green (control) and yellow (ΔN-β-catenin) arrow lines indicate the diameter of the contracted gels after

72 hours.

(K) Statistical quantification of gel contraction (% of initial area) by control and ΔN-β-catenin-overexpressing fibroblasts after a 72-hour incubation. The data are presented as mean ± SD (n = 5 replicates per group).

(L) Representative CRM images of gels embedded with control or ΔN-β-catenin-overexpressing fibroblasts after a 72-hour incubation.

(M–N) CRM images were analyzed using ImageJ with the BoneJ plugin to compare the connectivity and spacing of collagen fiber networks in the gels embedded with control and ΔN-β-catenin-overexpressing fibroblasts at 72 hours (n = 11 randomly selected 40X fields per group).

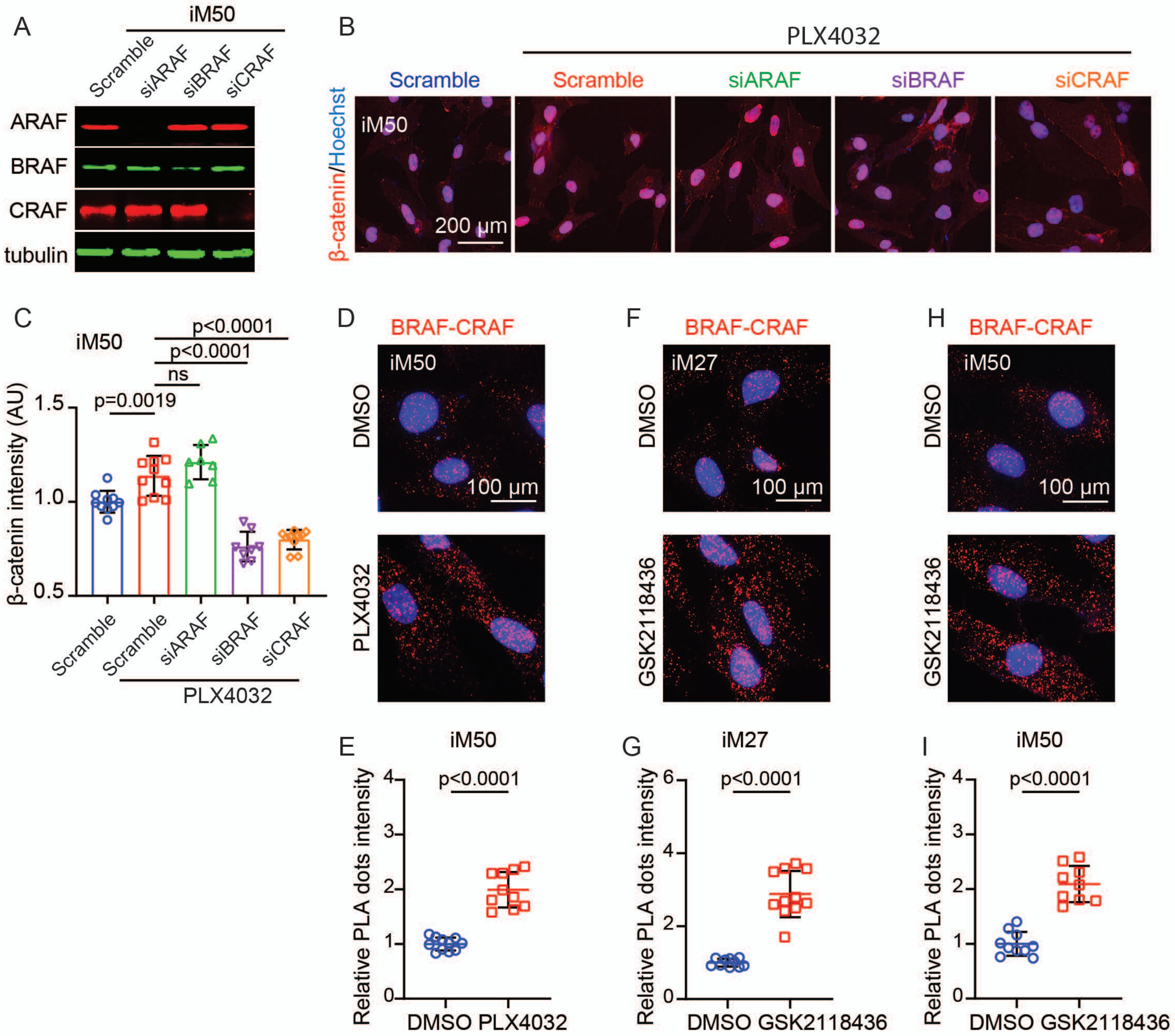

A
iM50
Scramble
siARAF
siBRAF
siCRAF
ARAF
BRAF
CRAF
tubulin
B
PLX4032
Scramble
Scramble
siARAF
siBRAF
siCRAF
β-catenin/Hoechst
iM50
200 μm
C
iM50
p<0.0001
p<0.0001
ns
p=0.0019
β-catenin intensity (AU)
1.5
1.0
0.5
Scramble
Scramble
siARAF
siBRAF
siCRAF
PLX4032
D
BRAF-CRAF
iM50
DMSO
100 μm
PLX4032
F
BRAF-CRAF
iM27
DMSO
100 μm
GSK2118436
H
BRAF-CRAF
iM50
DMSO
100 μm
GSK2118436
E
iM50
p<0.0001
Relative PLA dots intensity
4
3
2
1
0
DMSO
PLX4032
G
iM27
p<0.0001
Relative PLA dots intensity
6
4
2
0
DMSO
GSK2118436
I
iM50
p<0.0001
Relative PLA dots intensity
4
3
2
1
0
DMSO
GSK2118436


**Fig. S4**

**Figure S4 BRAFis bind to the RAF kinase domain and promote BRAF and CRAF dimerization.**

(A) Western blot confirming effective silencing of ARAF, BRAF, or CRAF expression in iM50 cells using corresponding siRNAs.

(B) Representative fluorescence images showing nuclear β-catenin expression in iM50 cells transfected with scramble siRNA or siRNAs that deplete ARAF, BRAF, or CRAF expression under PLX4032 treatment. iM50 cells transfected with scramble siRNA without PLX4032 treatment were used as a control.

(C) Quantification of nuclear β-catenin intensity in iM50 cells and genetically modified iM50 cells as shown in (B) with or without PLX4032 treatment using ImageJ. n = 8 randomly selected 20 X fields per group.

(D, F, H) Representative PLA images showing BRAF-CRAF heterodimerization in iM50 cells treated with DMSO or PLX4032 (D), in iM27 cells treated with DMSO or GSK2118436 (F), and in iM50 cells treated with DMSO or GSK2118436 (H). Red dots indicate BRAF-CRAF dimers.

(E, G, I) Quantification of PLA signals corresponding to panels (D), (F), and (H), respectively. Data are presented as mean ± SD (n = 9-11 randomly selected 40X fields per group).

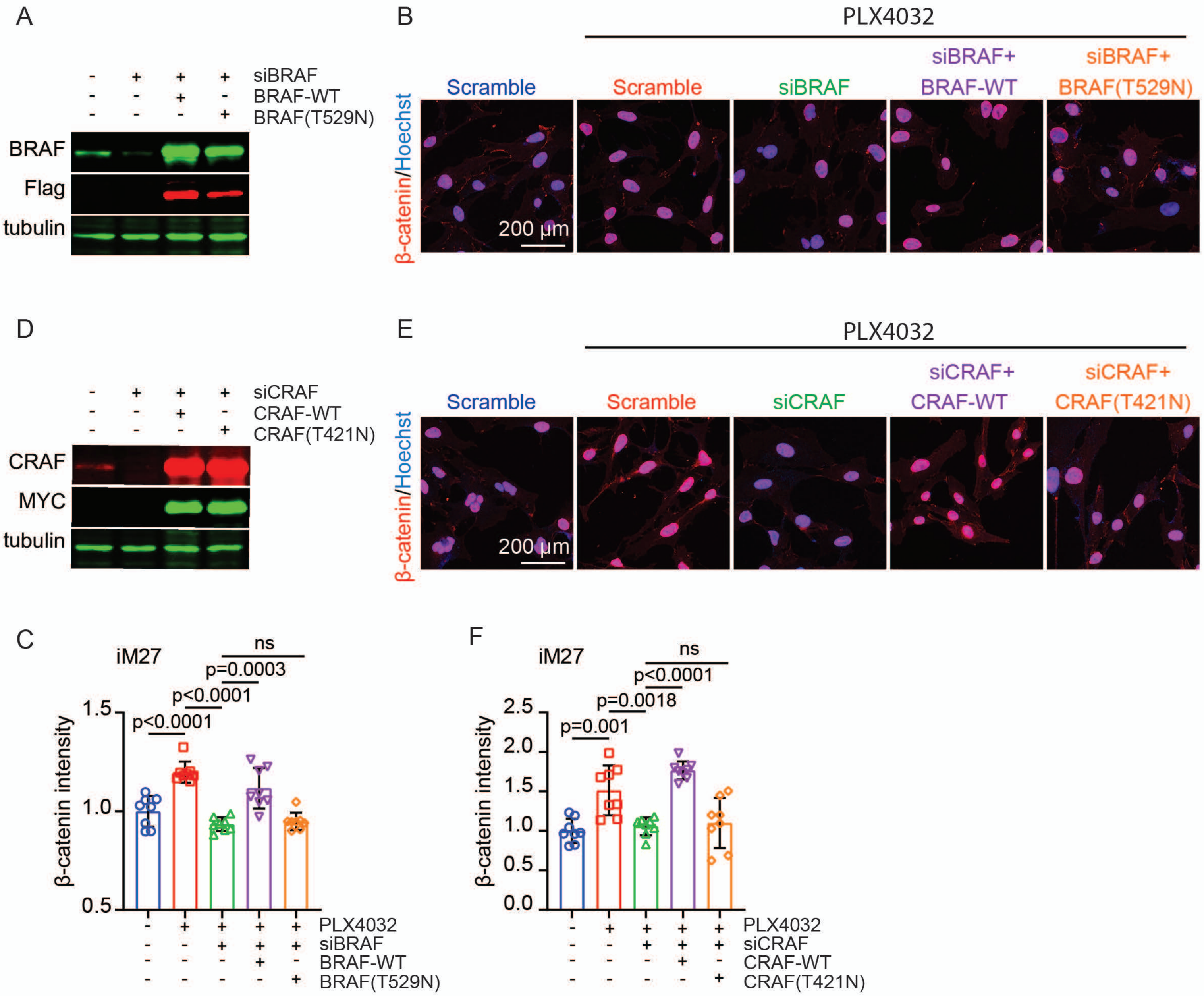

A
siBRAF
BRAF-WT
BRAF(T529N)
BRAF
Flag
tubulin
B
PLX4032
Scramble
Scramble
siBRAF
siBRAF+ BRAF-WT
siBRAF+ BRAF(T529N)
β-catenin/Hoechst
200 μm
D
siCRAF
CRAF-WT
CRAF(T421N)
CRAF
MYC
tubulin
E
PLX4032
Scramble
Scramble
siCRAF
siCRAF+ CRAF-WT
siCRAF+ CRAF(T421N)
β-catenin/Hoechst
200 μm
C
iM27
ns
p=0.0003
p<0.0001
p<0.0001
β-catenin intensity
PLX4032
siBRAF
BRAF-WT
BRAF(T529N)
F
iM27
ns
p<0.0001
p=0.0018
p=0.001
β-catenin intensity
PLX4032
siCRAF
CRAF-WT
CRAF(T421N)


**Fig. S5**

**Figure S5 BRAF and CRAF kinase domains are involved in PLX4032-induced nuclear β-catenin accumulation.**

(A) Western blot showing BRAF expression in iM27 cells transfected with scramble siRNA, iM27 cells transfected with BRAF siRNA (BRAF-deficient iM27), BRAF-deficient iM27 cells overexpressing wild-type BRAF (siBRAF + BRAF-WT), and BRAF-deficient iM27 cells overexpressing BRAF (T529N) (siBRAF + BRAF (T529N)).

(B) Representative fluorescence images showing nuclear β-catenin expression in iM27 and genetically modified iM27 cells shown in (A) with or without PLX4032 treatment as indicated.

(C) Quantification of the nuclear β-catenin intensity in iM27 cells and genetically modified iM27 cells shown in (B). n = 8 randomly selected 20X fields per group.

(D) Western blot showing CRAF expression in iM27 cells transfected with scramble siRNA, iM27 cells transfected with CRAF siRNA (CRAF-deficient iM27), CRAF-deficient iM27 cells overexpressing wild-type CRAF (siCRAF + CRAF-WT), and CRAF-deficient iM27 cells overexpressing CRAF (T421N) (siCRAF + CRAF (T421N)).

(E) Representative fluorescence images showing nuclear β-catenin expression in iM27 cells and genetically modified iM27 cells shown in (D) with or without PLX4032 treatment as indicated.

(F) Quantification of the nuclear β-catenin intensity in iM27 cells and genetically modified iM27 cells shown in (E). n = 8 randomly selected 20X fields per group.

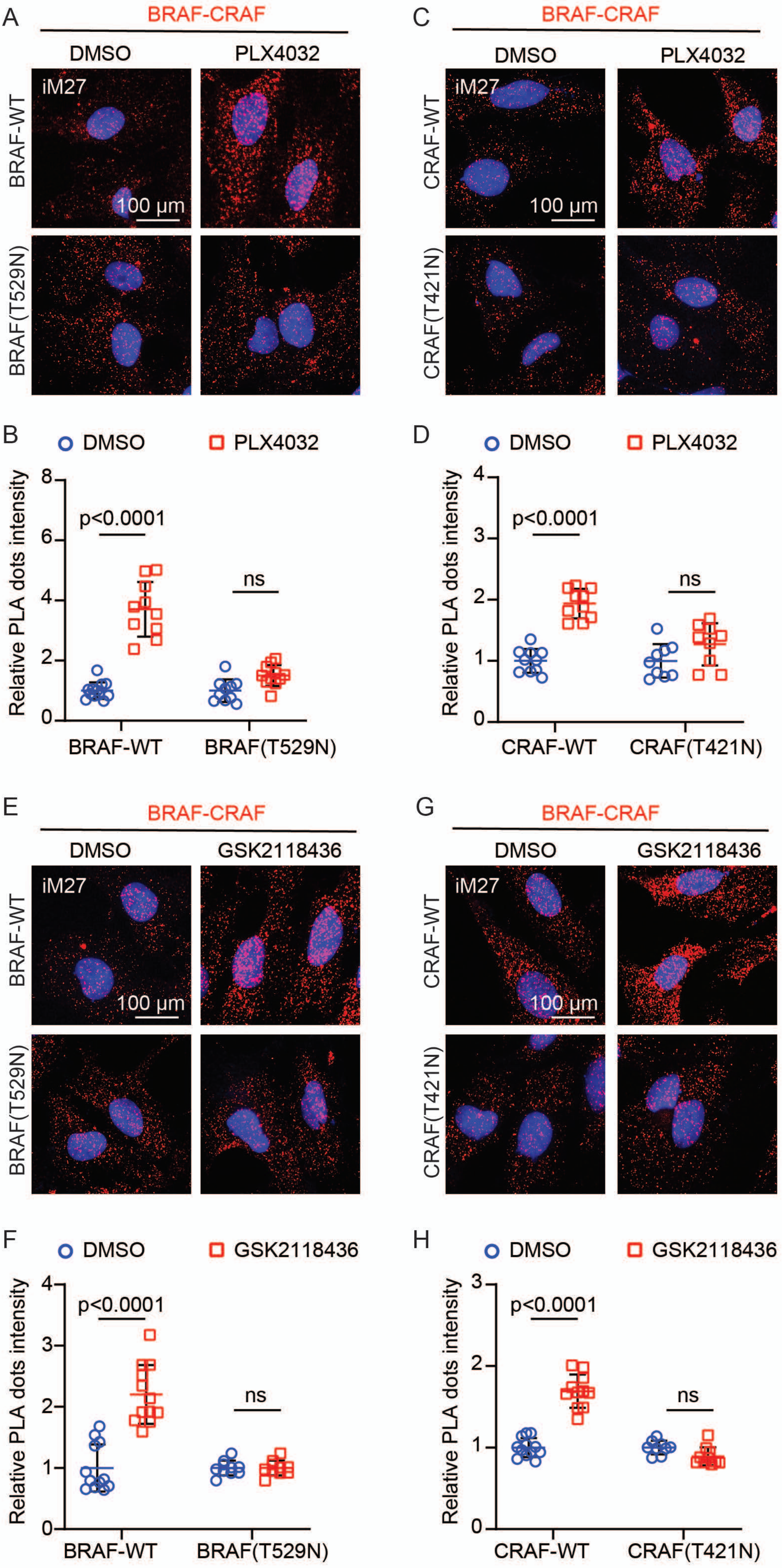
A
BRAF-CRAF
DMSO
PLX4032
iM27
BRAF-WT
BRAF(T529N)
100 µm
B
DMSO
PLX4032
Relative PLA dots intensity
p<0.0001
ns
BRAF-WT
BRAF(T529N)
C
BRAF-CRAF
DMSO
PLX4032
iM27
CRAF-WT
CRAF(T421N)
100 µm
D
DMSO
PLX4032
Relative PLA dots intensity
p<0.0001
ns
CRAF-WT
CRAF(T421N)
E
BRAF-CRAF
DMSO
GSK2118436
iM27
BRAF-WT
BRAF(T529N)
100 µm
F
DMSO
GSK2118436
Relative PLA dots intensity
p<0.0001
ns
BRAF-WT
BRAF(T529N)
G
BRAF-CRAF
DMSO
GSK2118436
iM27
CRAF-WT
CRAF(T421N)
100 µm
H
DMSO
GSK2118436
Relative PLA dots intensity
p<0.0001
ns
CRAF-WT
CRAF(T421N)

**Fig. S6**

**Figure S6 BRAF and CRAF kinase domains are involved in BRAFi-induced BRAF and CRAF dimerization.**

(A, C) Representative PLA images showing BRAF-CRAF heterodimerization in BRAF-deficient iM27 cells expressing wild-type BRAF or BRAF (T529N) (A) and in CRAF-deficient iM27 cells expressing wild-type CRAF or CRAF (T421N) (C). The cells were treated with PLX4032 and compared with the DMSO-treated controls. The red dots correspond to BRAF-CRAF interactions. (B, D) Quantification of the PLA signals (red dots) corresponding to panels (A) and (C). The data are presented as mean ± SD (n = 9-12 randomly selected 40X fields per group).

(E, G) Representative PLA images showing BRAF-CRAF heterodimerization in BRAF-deficient iM27 cells expressing wild-type BRAF or BRAF (T529N) and treated with DMSO or GSK2118436 (E), and in CRAF-deficient iM27 cells expressing wild-type CRAF or CRAF (T421N) and treated with DMSO or GSK2118436 (G). The red dots indicate BRAF-CRAF dimers. (F, H) Quantification of the PLA signals corresponding to panels (E) and (G). The data are presented as mean ± SD (n = 9-12 randomly selected 40X fields per group).

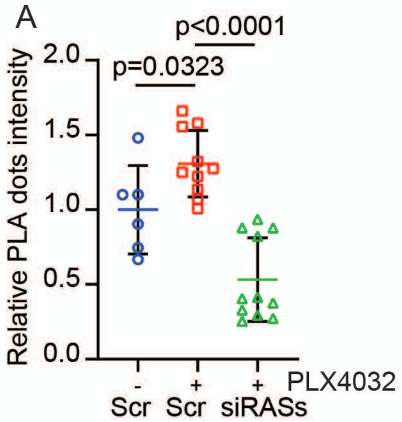

A
p<0.0001
p=0.0323
Relative PLA dots intensity
2.0
1.5
1.0
0.5
0.0
-
+
+
PLX4032
Scr
Scr
siRASs


**Fig. S7**

**Figure S7 RAS is involved in PLX4032-induced BRAF and CRAF heterodimerization**

(A) Quantification of the PLA signals (red dots) shown in Fig. 6B. The data are presented as mean ± SD (n = 6-11 randomly selected 40X fields per group).

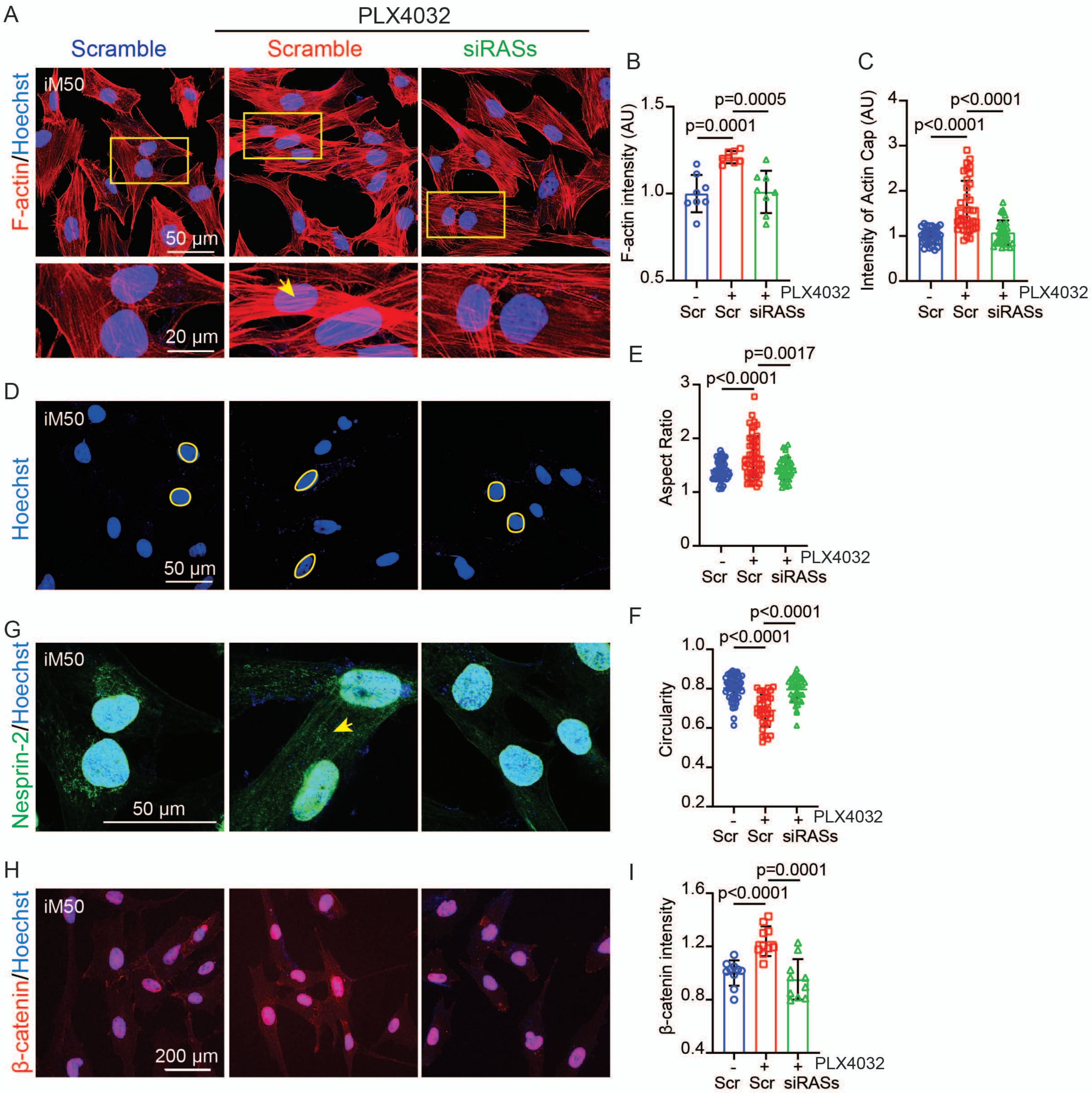

A
PLX4032
Scramble
Scramble
siRASs
F-actin/Hoechst
iM50
50 μm
20 μm
B
F-actin intensity (AU)
p=0.0005
p=0.0001
1.5
1.0
0.5
- + + PLX4032
Scr Scr siRASs
C
Intensity of Actin Cap (AU)
p<0.0001
p<0.0001
4
3
2
1
0
- + + PLX4032
Scr Scr siRASs
D
Hoechst
iM50
50 μm
E
Aspect Ratio
p=0.0017
p<0.0001
3
2
1
0
- + + PLX4032
Scr Scr siRASs
G
Nesprin-2/Hoechst
iM50
50 μm
F
Circularity
p<0.0001
p<0.0001
1.0
0.8
0.6
0.4
0.2
- + + PLX4032
Scr Scr siRASs
H
β-catenin/Hoechst
iM50
200 μm
I
β-catenin intensity
p=0.0001
p<0.0001
1.6
1.2
0.8
0.4
- + + PLX4032
Scr Scr siRASs


**Fig. S8**

**Figure S8 BRAFi-induced nuclear deformation and β-catenin accumulation are RAS dependent**

(A) Confocal images of F-actin expression and organization in iM50 transfected with scramble siRNA, scramble iM50 treaded with PLX4032 (scramble), and RAS-deficient iM50 (siRASs) treated with PLX4032. Insets display enlarged views of representative individual cells (highlighted by yellow boxes). Yellow arrows indicate the actin caps.

(B–C) Quantification of F-actin intensity (B) and actin cap intensity (C) using ImageJ. (n = 8 randomly selected 20X fields per group for B; n = 34-35 nuclei per group for C).

(D) Representative confocal images of nuclei visualized by Hoechst staining in iM50 transfected with scramble siRNA, scramble iM50 treaded with PLX4032, and RAS-deficient iM50 (siRASs) treated with PLX4032. The nuclear boundaries are outlined with yellow circles.

(E–F) Quantification of nuclear morphology based on the confocal images shown in (D), including the nuclear aspect ratio (E) and circularity (F), was performed via ImageJ. The data are presented as mean ± SD (n = 35-52 nuclei per group).

(G) Confocal images showing Nesprin-2 distribution in iM50 transfected with scramble siRNA, scramble iM50 treaded with PLX4032, and RAS-deficient iM50 (siRASs) treated with PLX4032. The yellow arrow indicates abnormal cytosolic localization of Nesprin-2.

(H) Representative fluorescence images showing nuclear β-catenin staining in iM50 transfected with scramble siRNA, scramble iM50 treaded with PLX4032, and RAS-deficient iM50 (siRASs) treated with PLX4032.

(I) Quantification of nuclear β-catenin intensity under the indicated conditions shown in (H). The data are presented as mean ± SD. n = 10 randomly selected 20 X fields per group.

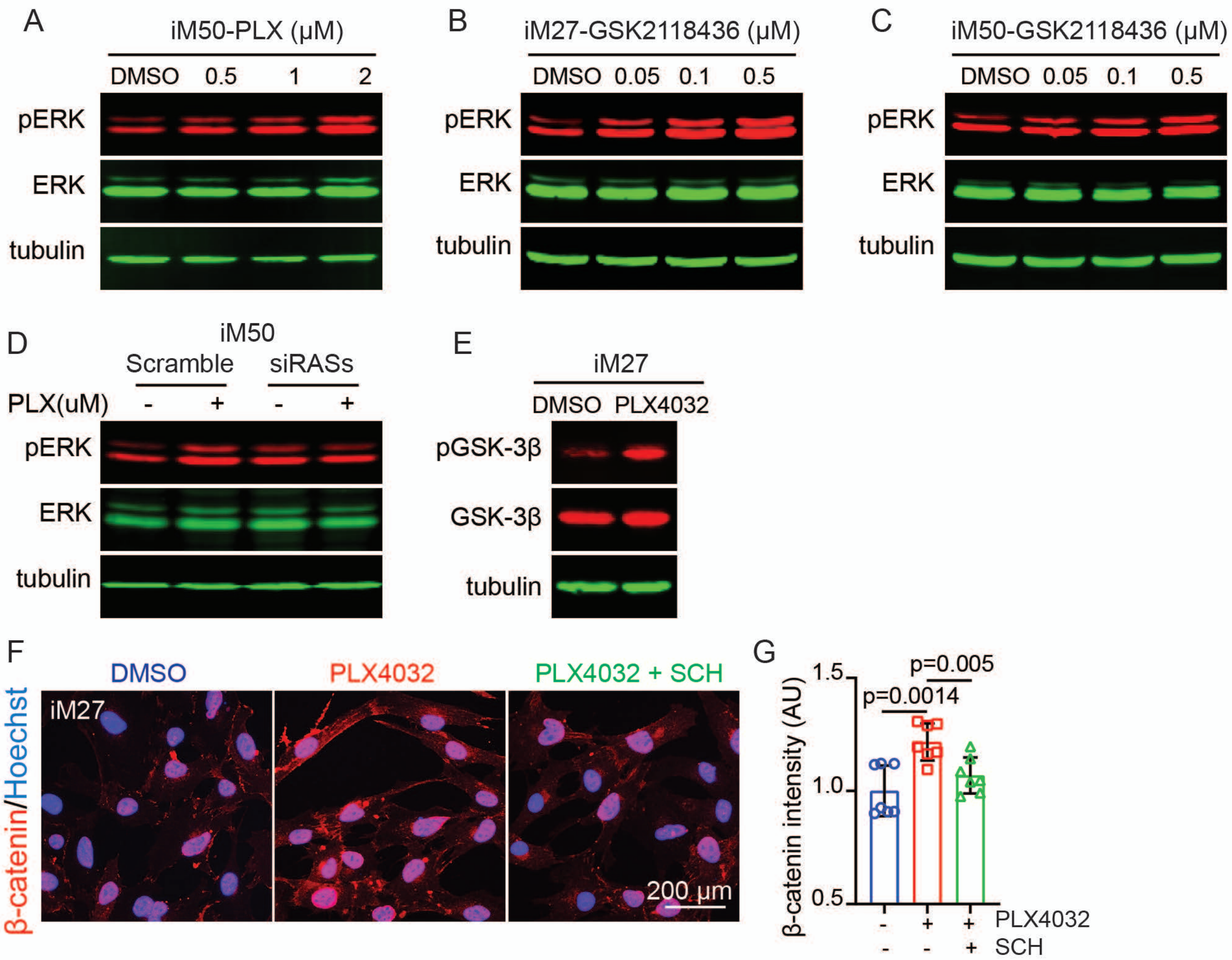
A
iM50-PLX (μM)
DMSO 0.5 1 2
pERK
ERK
tubulin
B
iM27-GSK2118436 (μM)
DMSO 0.05 0.1 0.5
pERK
ERK
tubulin
C
iM50-GSK2118436 (μM)
DMSO 0.05 0.1 0.5
pERK
ERK
tubulin
D
iM50
Scramble siRASs
PLX(uM) - + - +
pERK
ERK
tubulin
E
iM27
DMSO PLX4032
pGSK-3β
GSK-3β
tubulin
F
DMSO
PLX4032
PLX4032 + SCH
β-catenin/Hoechst
iM27
200 μm
G
β-catenin intensity (AU)
1.5
1.0
0.5
p=0.0014
p=0.005
PLX4032 - + +
SCH - - +

**Fig. S9**

**Fig. S9 ERK signaling is downstream BRAFi-induced RAF activation**

(A) Western blot showing increased ERK phosphorylation in iM50 cells in response to increasing PLX4032 concentrations compared with that in DMSO-treated cells.

(B–C) Western blot results showing increased ERK phosphorylation in response to increasing concentrations of GSK2118436 in iM27 cells (B) and iM50 cells (C).

(D) Western blot showing ERK phosphorylation in scramble siRNA-transfected iM50 and Ras-deficient-iM50 cells (siRASs) with or without PLX4032 treatment.

(E) Western blot showing increased GSK-3β phosphorylation in iM27 cells with and without PLX4032 treatment.

(F) Fluorescence microscopy images showing nuclear β-catenin expression in iM27 cells treated with DMSO, PLX4032, or a combination of PLX4032 and the ERK inhibitor SCH772984.

(G) Quantification of the nuclear β-catenin intensity in iM27 cells corresponding to (F) as shown. The data are presented as mean ± SD (n = 7 randomly selected 20 X fields per group).

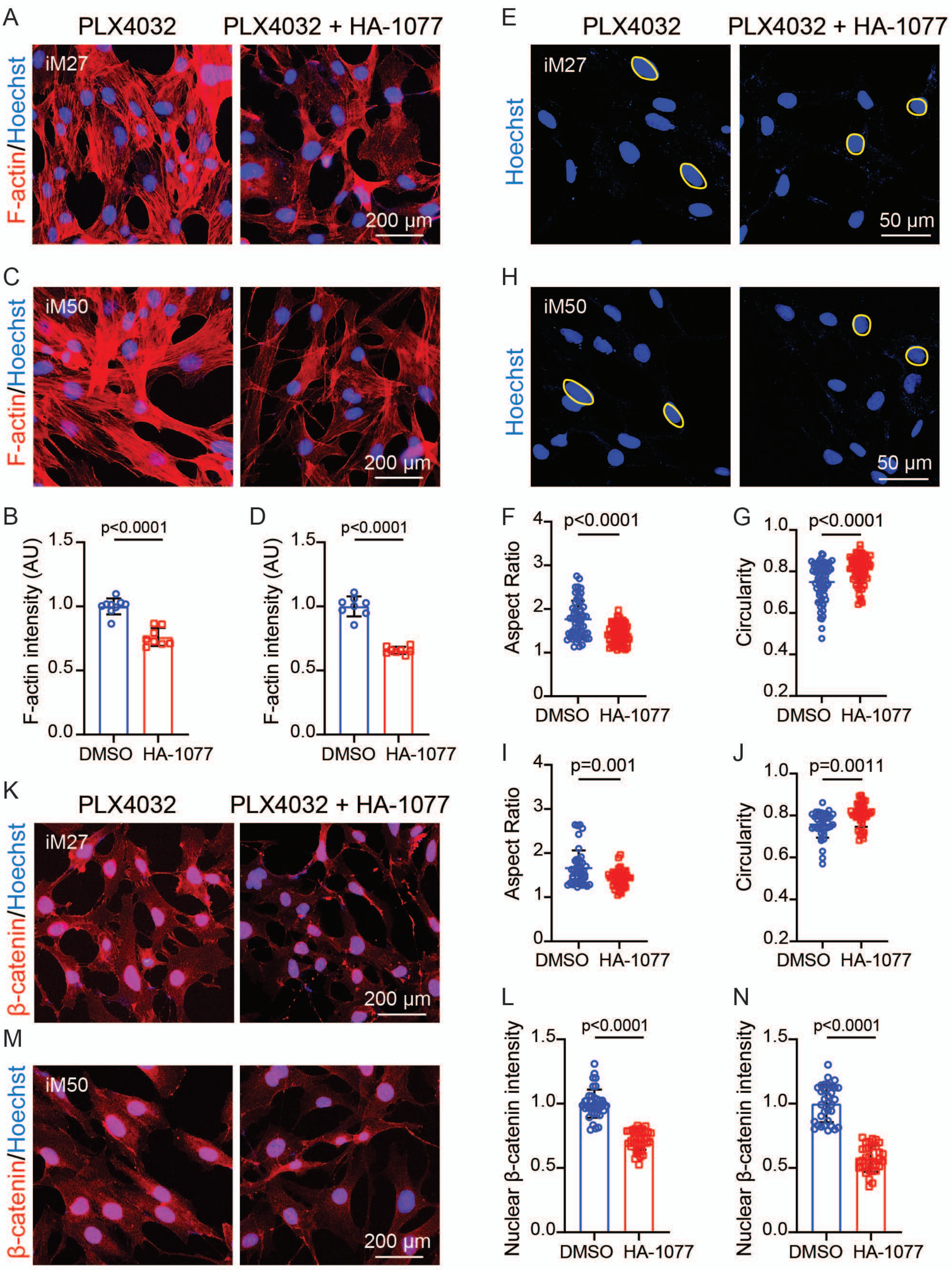
A
PLX4032
PLX4032 + HA-1077
F-actin/Hoechst
iM27
200 μm
C
iM50
200 μm
B
p<0.0001
F-actin intensity (AU)
DMSO
HA-1077
D
p<0.0001
F-actin intensity (AU)
DMSO
HA-1077
E
PLX4032
PLX4032 + HA-1077
Hoechst
iM27
50 μm
H
iM50
50 μm
F
p<0.0001
Aspect Ratio
G
p<0.0001
Circularity
I
p=0.001
Aspect Ratio
J
p=0.0011
Circularity
K
PLX4032
PLX4032 + HA-1077
β-catenin/Hoechst
iM27
200 μm
M
iM50
200 μm
L
p<0.0001
Nuclear β-catenin intensity
N
p<0.0001
Nuclear β-catenin intensity

**Fig. S10**

**Fig. S10 ROCK inhibition reverses PLX4032-induced nuclear deformation, actin polymerization, and β-catenin nuclear accumulation in CAFs.**

(A, C) Representative fluorescence images of F-actin staining in iM27 cells (A) and iM50 cells (C) treated with PLX4032 or a combination of PLX4032 and HA-1077.

(B, D) Quantification of the F-actin intensity in iM27 cells (B) and iM50 cells (D) under the indicated conditions corresponding to (A) and (C), respectively. The data are presented as mean ± SD (n=8-9 randomly selected 20X fields per group).

(E, H) Representative confocal images showing nuclear morphology visualized by Hoechst staining in iM27 cells (E) and in iM50 cells (H) treated with PLX4032 and a combination of PLX4032 and the ROCK inhibitor HA-1077.

(F–G) Quantification of nuclear morphology in iM27, including the nuclear aspect ratio (F) and circularity (G), was performed using ImageJ based on the corresponding images in (E). The data are presented as mean ± SD (n = 39-64 nuclei per group).

(I–J) Quantification of nuclear morphology in iM50, including the nuclear aspect ratio (I) and circularity (J), was performed using ImageJ based on the corresponding images in (H). The data are presented as mean ± SD (n = 39-64 nuclei per group).

(K, M) Representative fluorescence images showing nuclear β-catenin intensity in iM27 cells (K) and iM50 cells (M) treated with PLX4032 or a combination of PLX4032 and HA-1077.

(L, N) Quantification of the nuclear β-catenin intensity in iM27 cells (L) and iM50 cells (N) under the indicated conditions corresponding to (K) and.(M), respectively. The data are presented as mean ± SD (n = 33-40 nuclei per group).

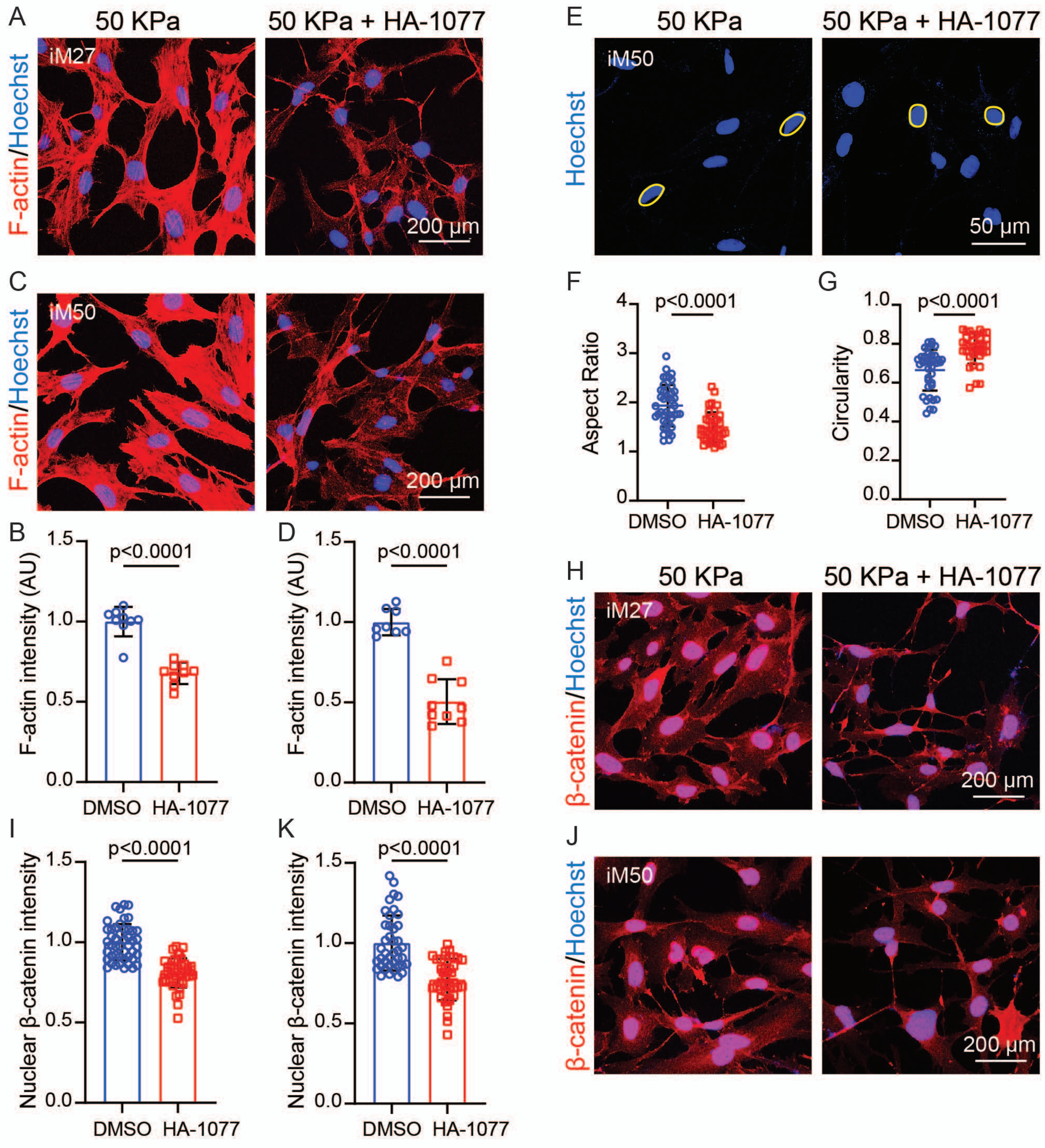
A
50 KPa
50 KPa + HA-1077
F-actin/Hoechst
iM27
200 μm
C
iM50
200 μm
B
D
F-actin intensity (AU)
p<0.0001
1.5
1.0
0.5
0.0
DMSO
HA-1077
E
Hoechst
iM50
50 μm
F
Aspect Ratio
G
Circularity
H
β-catenin/Hoechst
J
I
K
Nuclear β-catenin intensity

Fig. S11

**Fig. S11 ROCK inhibition reverses stiffness-induced nuclear deformation, actin polymerization, and β-catenin nuclear accumulation in CAFs.**

(A, C) Fluorescence microscopy images showing F-actin staining in iM27 cells (A) and iM50 cells (C) cultured on hard slides with a stiffness of 50 kPa with or without HA-1077 treatment.

(B, D) Quantification of the F-actin intensity in iM27 cells (B) and iM50 cells (D) corresponding to (A) and (C), respectively. The data are presented as mean ± SD (n = 8-9 randomly selected 20 X fields per group).

(E) Representative confocal images of Hoechst-stained nuclei in iM50 cells cultured on hard slides with a stiffness of 50 kPa with or without ROCK inhibitor HA-1077 treatment. Yellow circles highlight the approximate nuclear boundaries.

(F–G) Scatter dot plots showing the nuclear morphological changes under the indicated conditions corresponding to (E). The nuclear aspect ratio (F) and circularity (G) were analyzed and quantified using ImageJ. The data are presented as mean ± SD (n = 35-43 nuclei per group).

(H, J) Fluorescence microscopy images showing nuclear β-catenin expression in iM27 cells (H) and iM50 cells (J) cultured on hard slides with a stiffness of 50 kPa with or without HA-1077 treatment.

(I, K) Quantification of the nuclear β-catenin intensity iM27 cells (I) and iM50 cells (K) corresponding to (H) and (J), respectively. The data are presented as mean ± SD (n = 40-48 cells per group).
